# *In vivo* modelling of patient genetic heterogeneity identifies concurrent Wnt and PI3K activity as a potent driver of invasive cholangiocarcinoma growth

**DOI:** 10.1101/2021.07.05.449722

**Authors:** Nicholas T. Younger, Mollie L. Wilson, Edward J. Jarman, Alison M. Meynert, Graeme R Grimes, Konstantinos Gournopanos, Scott H. Waddell, Peter A. Tennant, David H. Wilson, Rachel V. Guest, Stephen J. Wigmore, Juan Carlos Acosta, Timothy J. Kendall, Martin S. Taylor, Duncan Sproul, Pleasantine Mill, Luke Boulter

**Author notes:** Corresponding Author: Dr Luke Boulter, MRC Human Genetics Unit, Institute of Genetics and Cancer, University of Edinburgh, EH4 2XU. both authors contributed equally to this manuscript (Nicholas T. Younger, Mollie L. Wilson). Centre for Inflammation Research, University of Edinburgh, Edinburgh, EH16 4TJ (Nicholas T. Younger, David H. Wilson).

## Abstract

Intrahepatic cholangiocarcinoma (ICC) is an aggressive and lethal malignancy of the bile ducts within the liver characterised by high levels of genetic heterogeneity. In the context of such genetic variability, determining which oncogenic mutations drive ICC growth has been difficult and developing modes of patient stratification and targeted therapies remains challenging. As a result, survival rates following a diagnosis with ICC have remained static since the late 1970s, whilst incidence of ICC has increased. Here, we performed the first functional *in vivo* study into the role that genetic heterogeneity plays in drivinga ICC via modelling of interactions between rare mutations with more common driver genes. By leveraging human ICC sequencing data to stratify and then model genetic heterogeneity in the mouse, we uncovered numerous novel tumour suppressors which, when lost, cooperate with the RAS oncoprotein to drive ICC growth. In this study, we specifically focus on a set of driver mutations that interact with KRAS to initiate aggressive, sarcomatoid-type ICC. We show that tumour growth of this cancer relies on both Wnt and PI3K signalling to drive proliferation and suppress apoptosis. Finally, we demonstrate that pharmacological co-inhibition of Wnt and PI3K *in vivo* substantially impedes the growth of ICC, regardless of mutational profile. As such, Wnt and PI3K activity should be considered as a signature by which patients can be stratified for treatment and inhibitors of these pathways should be levied as a treatment for patients diagnosed with ICC.

## Introduction

Intrahepatic cholangiocarcinomas (ICC) are epithelial tumours of the bile duct and are comprised of malignant ducts surrounded by an extensive stroma(1). ICC driven by infection with the liver fluke, *Opisthorchis viverrini* is endemic in South East Asia and whilst historically seen as a rare malignancy in the West, sporadic, non-fluke associated disease has increased in incidence in the UK, Europe, and the USA over the last four decades(2). In South East Asia ICC represents a significant health burden resulting in substantial social-economic impact(3). Currently, surgical resection is the only curative option for patients diagnosed with this cancer; however, of the ∼30% of patients who have disease that is amenable to surgery, 70% of those patients relapse following resection(4). In patients where surgery is not an option, the standard of care is palliative chemotherapy, which extends life by ∼3-6 months(5). Early studies using either patient ICC samples(6,7) or mouse models(8–10) demonstrated that oncogenic mutations in *Kras* (typically *Kras*^G12D^) and loss of function mutations in *Trp53* cooperate to initiate tumour formation. Recent genomic data, however, has challenged whether mutations in this oncogene and tumour suppressor pair often co-occur in human ICC(11,12). Instead, these sequencing data suggest that alternate or less-frequent mutations cooperate with more dominant oncogenes (such as mutant *Kras*) to promote tumorigenesis. Deep sequencing of ICC has uncovered a high level of genetic heterogeneity exists within patient ICC samples(13–15). Whilst a recurring set of mutations in canonical genes has been identified(1), many infrequent mutations have also been detected. However, the functional contribution of these infrequent *de novo* changes to affect disease progression and modulate therapeutic resistance or susceptibility remains unclear.

In order to identify and prioritise gain of function or loss of function mutations in a patient dataset of ICC, we use a computational pipeline, IntOGen(16), to generate a high-confidence list of candidate driver genes, of which 64 have not previously been assigned as being cancer drivers. To recapitulate the clonal competition observed in human tumorigenesis, we developed an *in vivo* CRISPR-SpCas9 system which simultaneously screens the candidate gene set against either *KRAS*^*G12D*^ or *NRAS*^*G12V*^ oncogenes. This identified a subset of genes in which human ICC-derived mutations genetically interact with RAS to initiate and accelerate ICC formation. Amongst these, we found that loss-of-function of Neurofibromin 2 (*Nf2*) interacts with mutant RAS to initiate tumour formation independent of *Trp53* status, highlighting again that as seen in patient data, RAS mutant cells do not strictly rely on *Trp53*-loss to initiate ICC. Loss of *Nf2* results in the formation of aggressive and poorly differentiated sarcomatoid-type ICC. These tumors are driven by dysregulation of Wnt-PI3K signalling, highlighting a novel therapeutic avenue which could be used to target ICC growth.

## Results

### Identifying candidate causative mutations that drive ICC growth

Intrahepatic cholangiocarcinoma contains a range of infrequently mutated genes without known function. Identifying a consensus group of driver mutations in ICC using exome and genome sequencing has been challenging, due in a large part to tissue availability. Nonetheless, a number of studies have demonstrated recurrent ICC mutations including neomorphic alterations in *IDH1* and *IDH2*, loss-of-function mutations in *PBRM1, BAP1, TP53, ARID1A* and gain-of-function mutations in *KRAS*. Despite their identification, the presence of these mutations in a tumour is not a strong predictor of therapeutic outcome(17–19) and for approximately 30% of ICC patients a driver mutation cannot be identified(20,21).

To determine whether all patient tumours contain potential driver mutations, we used a computational pipeline based around the driver prediction tool IntOGen(16). This method utilises a combination of functional impact bias (OncodriveFM), spatial clustering (OncodriveCLUST) and corrected frequency (MutSigCV) to define whether particular genomic regions have a mutational rate beyond that which is expected, have a bias towards clustered mutations or those which are likely to impact functional domains, such as those that are regulatory or catalytic (summarised in **Supplementary Figure 1A**). Having filtered out hypermutated samples (**Supplementary Table 1**), we used this pipeline to analyse the variants identified in the genomes of 277 sporadic and fluke-associated ICCs from four distinct studies(12,14,22–24), summary information on the aggregated cohort can be found in **Supplementary Figure 2A-H** and **Supplementary Table 2**). Following processing, 55% of samples (N=152) carried ≥2 predicted drivers whilst 18% (N=50) carried none (**Figure 1A**, and **Supplementary Table 3**). Of these predicted driver mutations, approximately one third of mutations were already known to occur in ICC or were present in genes in the COSMIC database (**Figure 1B**). The remaining two thirds of mutations were novel and occurred in fewer than 8% of ICC cases. Indeed, the majority of novel mutations were only found in 3-4 patients, corresponding to ∼1.5% of the patient cohort (**Figure 1C, Supplementary Tables 4** and **5**). To explore whether these low-frequency predicted drivers are involved in common pathways or processes, networks were constructed based on known and predicted physical interactions(25,26) and clustered into modules based on connection density(27) (**Figure 1D, Supplementary Table 6**). This produced a network with 6 modules containing a mix of known in ICC, COSMIC, and novel genes. Gene ontology analysis was performed on the modules to ascertain the biological processes in which each module may participate (**Supplemental Table 6**). Processes featuring isolated novel drivers or novel drivers in combination with better-known cancer genes included negative regulation of cell growth (*APBB1, STK11*), negative regulation of cell proliferation and migration (*NF2*), and regulation of angiogenesis (*EPHA2*). Importantly, the connectivity between novel and known genes in shared processes suggests that whilst mutated at low frequency in the patient population, these novel drivers may contribute to cancer phenotypes by disrupting common processes associated with more established cancer genes.

**Figure 1.**
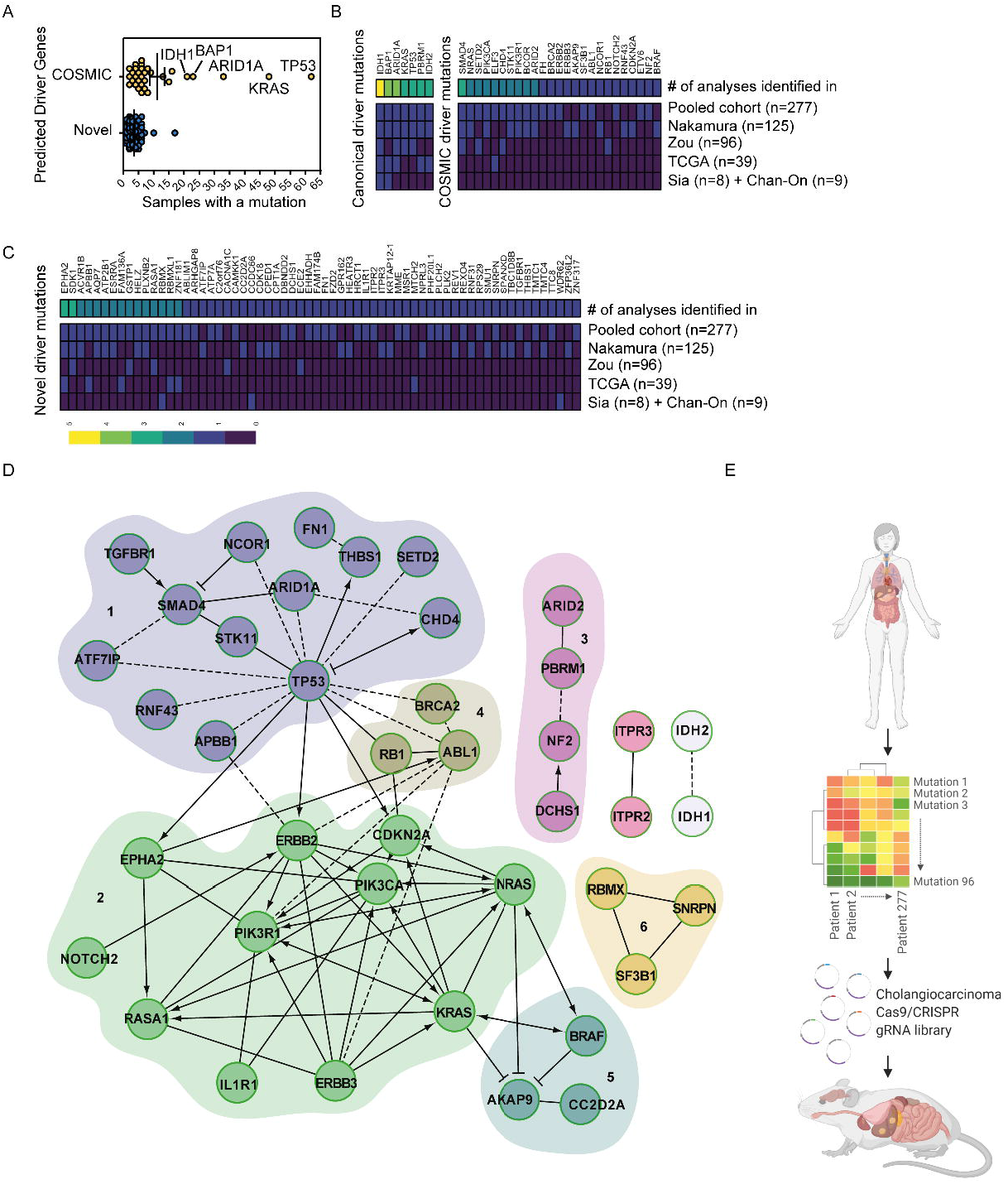
*In silico* screening identifies novel drivers of intrahepatic cholangiocarcinoma: **A**. The number of samples with mutations in driver genes identified following analysis with IntOGen. Samples clustered into those which had been previously identified in ICC or are present in the COSMIC database (yellow points), or those mutated genes that have not previously been assigned as being cancer drivers (blue points). **B**. Frequency of samples containing known ICC or COSMIC mutation in each of the individual cohorts collated in this study and in the pooled datasets and **C**. Frequency of samples containing a predicted novel oncogenic mutation called by IntOGen. In both C and D heat maps represent the frequency at which each mutation is found within each study and the aggregated frequency between all studies. Upper bar represents the number of times a mutation was identified between studies. **D**. Pathway interaction analysis of putative ICC driver mutations identified by IntOGen based on known functional (solid lines) and predicted physical (dotted lines) interactions. Numbers 1-6 represent distinct gene-relationship modules based on predicted or known genetic interactions. **E**. Schematic study representation in which whole genome and whole exome sequencing has been collated from patients with ICC. The mutational profile of these tumours was rationalised to identify novel, high confidence drivers of ICC. These putative drivers were used as input for an *in vivo* SpCas9/CRISPR screen, which generated tumours in the adult mouse liver. These were used to identify novel functional processes that drive ICC growth. N=277 patient exomes or genomes with matched, non-cancerous tissue.

### *In vivo* CRISPR-SpCas9 screening identifies novel tumour suppressors in RAS driven ICC

Clonal analysis of patient ICC has failed to identify a consensus mutational route through which tumours progress(28) or epistatic mutations that functionally interact to drive tumour initiation and growth. Relatively low sample number and high genetic heterogeneity in ICC exacerbate the difficulties with this type of associative analysis.

To overcome these limitations and define which candidate drivers are functionally capable of initiating ICC, we developed an *in vivo* screening approach that allowed us to functionally prioritise ICC driver mutations (**Figure 1E**). Previous work using multiplex-mutagenesis in the liver has demonstrated that editing specific genomic loci in hepatocytes can give rise to ICC, albeit using a relatively limited pool of gRNAs, targeting 10 genes(29). In this system, naked DNA is delivered to the liver using a high pressure, hydrodynamic injection into the lateral tail-vein of mice. When paired with CRISPR-SB plasmids containing sgRNAs and SpCas9 flanked between two Sleeping Beauty (SB) inverted terminal repeats, and a plasmid expressing Sleeping Beauty (SB) transposase, it is possible to edit *in vivo* endogenous genes in hepatocytes in a mosaic manner(30). Using the CRISPR-SB system as a starting point, we generated a large-scale multiplexed CRISPR-SpCas9 plasmid library (known hereafter as ICC^Lib^) containing triplicate gRNAs targeting 91 mouse homologues of our putative, patient-derived ICC driver genes identified through our *in silico* approach. Five genes (*KRTAP12-1, RBMXL1, RBMX, SPANXD* and *ZNF181*) from our human dataset had no identifiable murine orthologue, and so were excluded from further analysis (**Supplementary Table 1** and **Supplementary Figure 3**).

We randomly introduced CRISPR-SpCas9 targeted mutations into these candidate “patient-led” ICC genes in otherwise wild-type mice to determine which loss of function mutations are necessary for tumour initiation. The ICC^Lib^ alone failed to induce any tumours in mice after 10 weeks suggesting that within this timeframe these loss-of-function mutations are insufficient to initiate cancer. Gain-of-function mutations in *KRAS* and *NRAS* have been previously described in ICC(13,14) and through our IntOGen analysis, we similarly identified recurrent mutations in both of these genes (*KRAS* 18.05% and *NRAS* 2.88%, **Supplementary Table 4**). In experimental models, expression of mutant RAS in the adult liver is weakly oncogenic and normally insufficient to initiate ICC formation; rather, mutant cells undergo oncogene-induced senescence and are removed from the liver by immune clearance(31). Therefore, we co-expressed GFP tagged KRAS^G12D^ or NRAS^G12V^ with our ICC^Lib^ to determine whether any of the loss-of-function mutations introduced via the ICC^Lib^ synergise with mutant RAS to promote ICC initiation. Within 10 weeks, mice that received either KRAS^G12D^ or NRAS^G12V^ and ICC^Lib^ developed macroscopic and multifocal cancer; this was accelerated in KRAS^G12D^ mice, which developed symptomatic liver cancer in 8 weeks (**Figure 2A**). In those mice that developed cancer, multiple tumours formed per mouse (**Figure 2B**) which were histologically aggressive adenocarcinoma with a poorly differentiated cholangiocellular morphology. Importantly, these tumours express GFP (**Supplementary Figure 4A**), denoting that they continue to express the KRAS^G12D^ or NRAS^G12V^ constructs, and the cholangiocyte marker Keratin-19, which is constrained to the biliary epithelium in normal livers (**Supplementary Figure 4B**).

**Figure 2.**
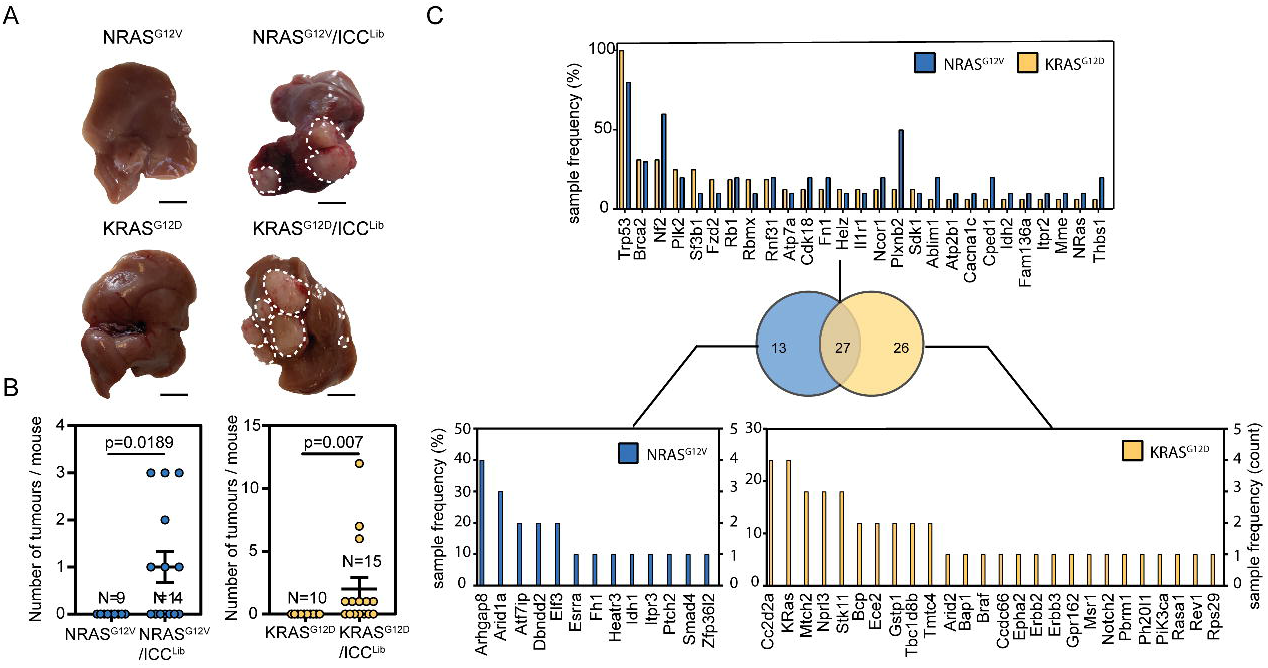
*In vivo* CRISPR-Cas9 screening identifies transforming mutations that interact with mutant Ras. **A**. Macroscopic images of the livers following injection with either NRAS^G12V^ or KRAS^G12D^ alone (left-hand images) or in combination with ICC^Lib^, right-hand images, dotted line denotes tumour (scale bar = 1 cm). **B**. Quantification of macroscopic tumours per mouse at 10 weeks in mice bearing Nras^G12V^ expressing tumours and 8 weeks in those with Kras^G12D^-driven cancer. Each circle represents a different animal. **C**. The number of samples containing Indels in a particular gene following whole exome sequencing. Upper graph lists those mutations found in both NRAS^G12V^ and KRAS^G12D^ tumours, and lower graphs denote those mutations that are found only in KRAS^G12D^ or NRAS^G12V^ expressing tumours. Sample frequency (%) denotes the proportion of tumours containing any given mutation, whereas (count) is absolute number. (N represents anatomically discreet tumours recovered from at least four individual animals, KRAS^G12D^ N=14 and NRAS^G12V^ N=10).

To determine which CRISPR-SpCas9 mutational events cooperated with KRAS^G12D^ or NRAS^G12V^ and lead to the emergence of liver cancer, whole exomes were sequenced from 14 KRAS^G12D^ and 10 NRAS^G12V^ driven tumours. All indels within 50bp of a sgRNA target site were manually inspected to determine whether they were Cas9-induced or of spontaneous origin. Almost all indels had start or end positions approximately 3bp upstream of the SpCas9 protospacer adjacent motif (PAM) sequence strongly indicating that they are a consequence of CRISPR-SpCas9 editing (**Supplementary Figure 5A**). Tumours acquired multiple CRISPR-SpCas9-induced lesions; KRAS^G12D^ tumours contained an average of 7.5±1.19 mutations and in NRAS^G12V^ tumours there were on average 7.7±2.59 CRISPR-SpCas9 induced mutations (**Supplementary Figure 5B**). Across both Kras^G12D^ and Nras^G12V^ screens, 66 of the 91 predicted drivers targeted with the ICC^Lib^ were mutated and 27 of these were shared between NRAS^G12V^ and KRAS^G12D^ tumours (**Figure 2C** and **Supplementary Figure 6A**). The most common CRISPR-SpCas9 mutation we identified was unsurprisingly in *Trp53*, reiterating the ability of cells with *Trp53*-loss-of-function mutations to overcome RAS-induced senescence(32). We also found recurrent CRISPR-induced indels from both NRAS^G12V^ and KRAS^G12D^ screens in genes whose loss has been linked to ICC, but which have not previously been shown to genetically interact with mutant RAS in this cancer including *Brca2, Nf2* and *Plk2* (**Figure 2C**).

### *Nf2-*loss interacts with KRAS^G12D^ and *Trp53*-loss to promote sarcomatoid phenotypes in ICC

Our data demonstrate that loss of numerous genes mutated at low frequency in human ICC potentially interact with activating NRAS^G12V^ and KRAS^G12D^ mutations to promote ICC initiation *in vivo*. These data and those of others has demonstrated that *KRAS* mutations occur more frequently in ICC than those in *NRAS(15)*, therefore we prioritized validating loss-of-function mutations that genetically interact with KRAS^G12D^. RNA sequencing analysis of our ICC^Lib^ screen tumours showed that, on the whole, tumours transcriptionally clustered closely to each other by Principal Component Analysis (PCA) and were transcriptionally similar to tumours generated by expressing both KRAS^G12D^ and a shRNA targeting *Trp53* (**Supplementary Figure 7**). Furthermore, our screen tumours were transcriptionally distinct from normal bile ducts. However four tumours in our screen transcriptionally segregated from all other tumours; all contained SpCas9/CRISPR-induced mutations in both *Trp53* and *Nf2*. In fact, of the 14 cancers from our screen that we exome sequenced, four of the five containing *Nf2*-mutations segregated away from the main cluster (**Supplementary Figure 7**), suggesting that the addition of a mutation in *Nf2* can functionally cooperate with *Kras*^*G12D*^ and *Trp53* mutations and affects the phenotype of ICC. NF2 is also known as Merlin and has a well-defined role in the Hippo/LATS signalling pathway, where it negatively regulates pathway activation through the phosphorylation of *Mst1/2*; however, NF2/Merlin is also known to interact with a number of other signalling pathways including PI3K and Wnt signalling(33,34). We elected to investigate the genetic interaction of *Nf*2, *Trp53* and *Kras*^*G12D*^ further, by generating gRNAs to specifically target *Nf2* and *Trp53* (or a non-targeting control, gRNA^scrm^) which were then co-injected hydrodynamically with our KRAS^G12D^ expressing construct. *Trp53-*loss and *Nf2*-loss were both capable of overcoming the senescence inducing effects of Kras^G12D^ expression in the liver and mice developed lethal tumours within 8 weeks following injection (**Figure 3A**). In the presence of KRAS^G12D^, the singular deletion of either *Trp53* or *Nf2* resulted in large discrete tumours. However, dual loss of *Trp53* and *Nf2* resulted in cancers that were highly aggressive and invasive, which had a median survival of 14 days compared to 39 and 41 days in singular *Nf2*-deleted and *Trp53*-deleted tumours, respectively (**Figure 3A**). Tumours that lacked both *Trp53* and *Nf2* were highly diffuse and covered less liver area compared to both single gene deletions; however, the number of tumours that formed was significantly higher in *Trp53*;*Nf2* co-deleted tumours, suggesting that mutations in these two tumour suppressors may synergise in cancer and are not functionally redundant (**Figure 3B** and **3C**). Histopathologically, the *Trp53*;*Nf2* co-deleted cancers are highly invasive with pan-cytokeratin and CK19 immunopositive cells migrating throughout the liver (**Figure 3D**) and represent a model to study the biology of invasive, sarcomatoid ICCs which migrate along the ducts and invade the liver(35,36). Rather than those ICCs which are mass forming (and which have been previously modelled in mice), sarcomatoid ICC whilst rare, has a very poor prognosis with a survival of weeks to months following diagnosis(37,38).

**Figure 3.**
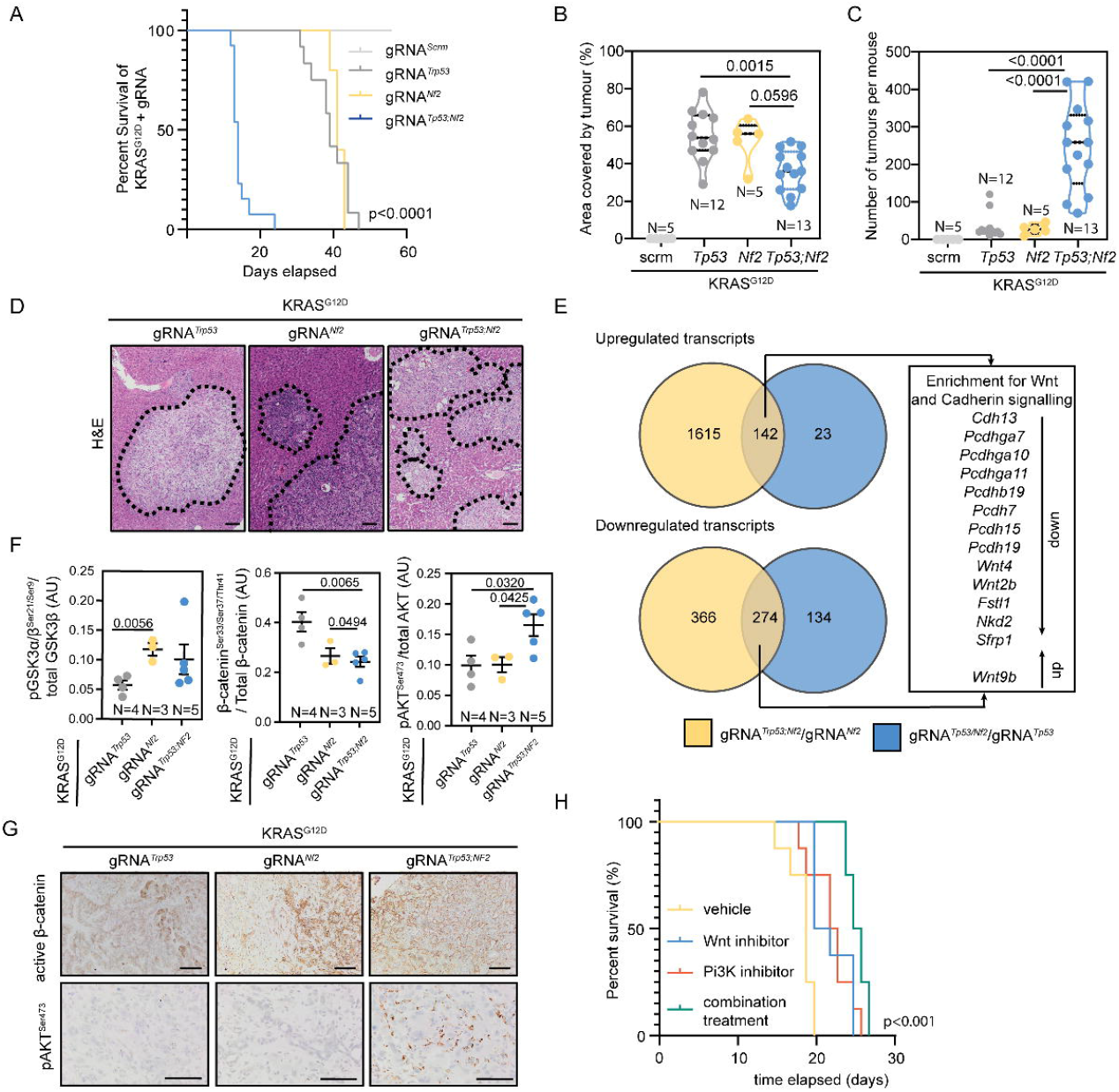
*Nf2*-loss results in Ras^G12D^-induced oncogenesis and cooperates with *Trp53*-loss to accelerate ICC formation. **A**. Kaplan-Meier curve demonstrating the relative survival proportions of mice with KRAS^G12D^ and gRNAs targeting *Trp53* (N=12), *Nf2* (N=5), *Nf2*;*Trp53* (N=13) or non-targeting control (scrm, N=5). **B**. Proportion of liver occupied by tumour and **C**. number of tumours per mouse. **D**. H&E staining of KRAS^G12D^ tumours with *Trp53, Nf2* or *Trp53* and *Nf2*-loss, scale bar = 200μm, dotted line denotes tumour boundary. **E**. Comparison of RNAseq analysis when the transcriptomes from *Nf2*;*Trp53* vs *Trp53*-alone tumours (blue) are compared to transcripts from *Nf2*;*Trp53* vs *Nf2*-alone (yellow) tumours. Each group contains N=4 regionally distinct tumours. **F**. Analysis of RPPA data demonstrating the changes in the proportion of phosphorylated GSK3, β-catenin and AKT relative to total protein levels in KRAS^G12D^;*Trp53*^KO^ (grey points), KRAS^G12D^;*Nf2*^KO^ (yellow points), KRAS^G12D^;*Trp53*^KO^;*Nf2*^KO^ (blue points). **G**. Immunohistochemistry of active, dephosphorylated β-catenin (upper panels) and phosphorylated AKT^Ser647^ (lower panels) in KRAS^G12D^;*Trp53*^KO^, KRAS^G12D^;*Nf2*^KO^, KRAS^G12D^;*Trp53*^KO^/*Nf2*^KO^ tumours. Scale bar = 50μm **H**. Kaplan-Meier curve demonstrating the survival changes when KRAS^G12D^;*Trp53*^KO^;*Nf2*^KO^ animals are treated with vehicle (yellow line), LGK974 (Wnt-inhibitor, blue line), Pictilisib (PI3K inhibitor, orange line) or a combination (green line). N=5 per group.

To determine the transcriptomic differences driving this aggressive phenotype when compared to individual *Trp53* or *Nf2* deleted tumours, we undertook bulk RNA sequencing of tumours with Kras^G12D^;*Trp53*^KO^, Kras^G12D^;*Nf2*^KO^ or Kras^G12D^;*Trp53*^KO^;*Nf2*^KO^ genetic profiles. Sarcomatoid, *Trp53*;*Nf2* co-deleted tumours are transcriptionally more distinct from cancers containing either *Trp53* or *Nf2* deletions alone based on PCA (**Supplementary Figure 8A** and **Supplementary Table 7**). By comparing the up- and down-regulated genes in *Trp53* vs *Trp53*;*Nf2* co-deleted tumours against the changes found in *Nf2* vs *Trp53*;*Nf2* co-deleted tumours, we identified enrichment in signatures for Wnt signalling and Cadherin signalling using PANTHER (**Supplementary Figure 8B**). However, we did not find a transcriptional signature for Hippo signalling, nor could we identify YAP positive cells within the *Nf2*-deleted tumours, rather YAP-positive cells are only found adjacent to the tumour mass, as previously described(39). Together, these data indicate that *Nf2*-loss in these tumours fails to activate Hippo signalling (**Figure 3E** and **Supplementary Figure 8B-C**). Previous work from our group has shown that Wnt signalling promotes ICC growth in the absence of classical Wnt pathway “activating mutations”(40). In *Trp53*;*Nf2* co-deleted cancers, we observe upregulation of *Wnt9b* and suppression of inhibitors of Wnt signalling *Nkd2* and *Sfrp2* suggesting that alterations in ligand levels and negative regulators of Wnt signalling are important mediators of ICC progression. Interestingly, sarcomatoid ICC in patients displays changes in cadherin expression, though it is not clear whether these changes are causative for the aggressive sarcomatoid phenotype(37). In our model, we found transcriptional suppression of a range of cadherins and proto-cadherins that have been also implicated in Wnt regulation(41,42).

As there are few targeted treatments for poorly differentiated ICC, we screened our models driven by KRAS^G12D^-expression and either *Tp53*-loss, *Nf2*-loss or a combination of the two for activated and pharmacologically targetable signalling pathways using Reverse Phase Protein Arrays (RPPAs). Deletion of *Nf2* results in increased inhibitory phosphorylation of GSK3β and activation of β-catenin signalling when compared to KRAS^G12D^ driven tumours lacking *Trp53* alone, suggesting that the ligand and inhibitor alterations we found at the transcriptional level translate into pathway activation (**Figure 3E** and **3F**). Furthermore, when *Trp53* and *Nf2* are concurrently deleted, the proportion of AKT that is phosphorylated at Serine-473 significantly increases (**Figure 3F**). Histologically, dephosphorylated, active β-catenin is found within KRAS-driven cancers (**Figure 3G**), however the accumulation of phosphorylated AKT^Ser473^ is highly restricted to cancers harbouring both *Trp53-* and *Nf2*-loss of function mutations. These data suggest that concurrent pAKT and Wnt activity promote the development of sarcomatoid ICC whilst suppressing apoptosis (**Supplementary Figure 9**) thereby promoting the rapid development of cancer in the liver. To test whether the aggressiveness of *Trp53*;*Nf2* mutated ICC is dependent on Wnt and AKT signalling we treated mice bearing KRAS^G12D^;*Nf2*;*Trp53*-KO tumours with an inhibitor of Porcupine (LGK974)(43), which reduces Wnt ligand secretion by preventing palmitoylation of Wnt ligands and a PI3K inhibitor, Pictilisib(44), which prevents the conversion of PIP_2_ into PIP_3_ and thereby reduces AKT phosphorylation. Both LGK974 and Pictilisib significantly improve survival of tumour bearing mice compared to animals treated with vehicle alone (median survival of 19 days in vehicle treated versus 21 days in LGK974-treated (p=0.0015) and 22.5 days in Pictilisib-treated animals (p=0.0058), respectively). Wnt inhibition and PI3K inhibition performed similarly, with no statistically significant difference in survival outcomes when used as single agents (**Figure 3H** and **Supplementary Figure 10A-C**). In combination, LGK974 and Pictilisib improve median survival from 19 days to 25.5 days (**Figure 3H**), significantly (p<0.001) reducing mortality compared to single treatments (**Supplementary Figure 10D-F**) and demonstrating that co-inhibition of Wnt and PI3K signalling is an effective treatment in sarcomatoid tumours that lack classical Wnt and PI3K activating mutations (i.e mutations in *APC, CTNNB1* and *PI3KCA*).

### Wnt and PI3K/AKT represent a conserved mechanism by which distinct pathological subtypes of ICC grow

From previous transcriptomic studies from ICC, it is possible to identify a subgroup of patients with high Notch pathway activity and who would be sensitive to treatment with γ-secretase-inhibitors(45). We therefore sought to identify whether there is a group of ICC patients who could be sensitive to co-inhibition of Wnt and PI3K signalling. In 104 transcriptomes from ICC patients(22), there is a high level of correlation (r=0.690, P<0.0001) between those with a high expression of genes associated with Wnt signalling and high expression of genes associated with AKT signalling (**Figure 4A** and **Supplementary Table 8**). Therefore, we considered whether the activation of Wnt and PI3K signalling is a more universal process in tumour formation and sought to address whether these pathways are recurrently activated in ICC lacking *RAS* mutations.

**Figure 4:**
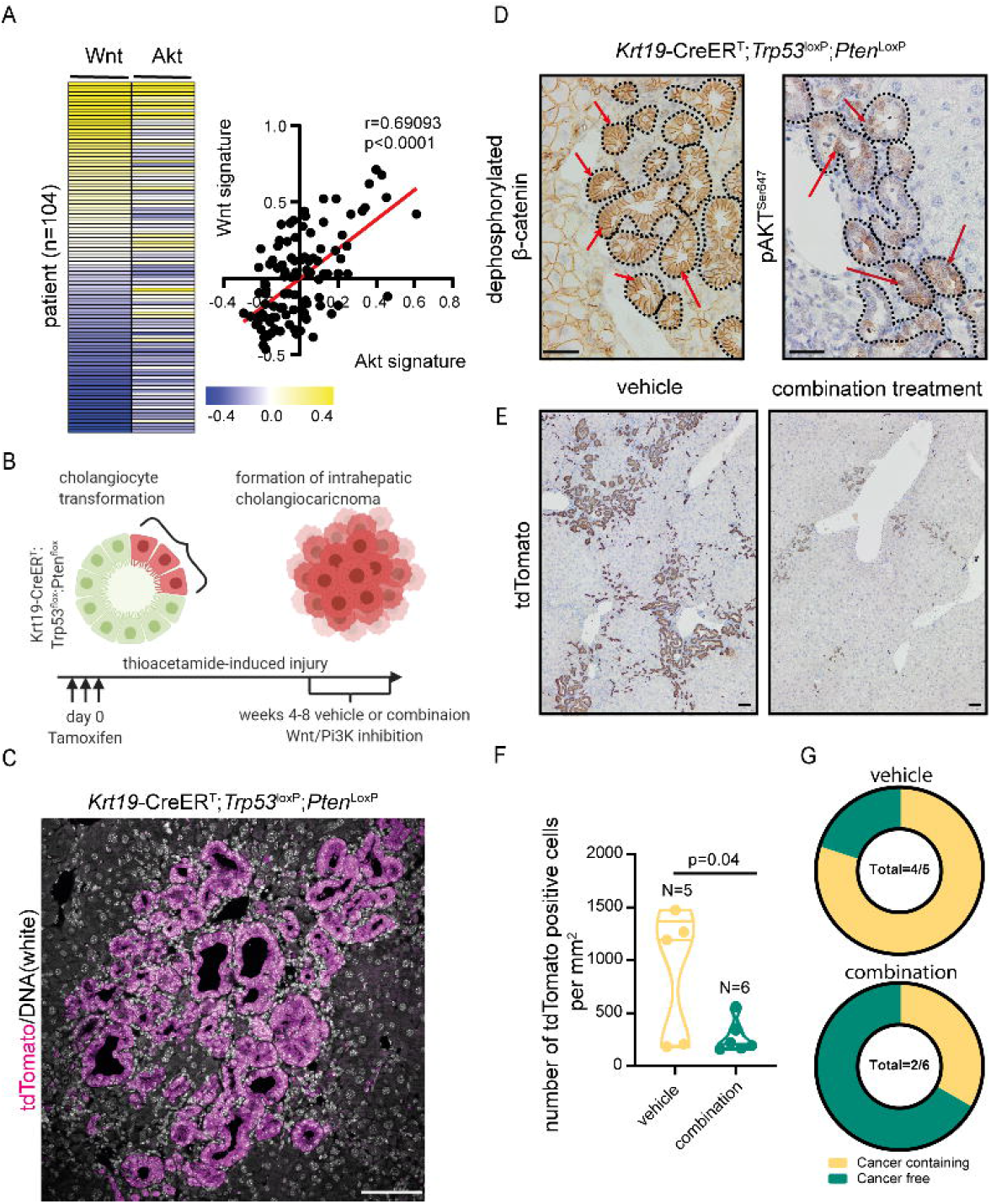
Therapeutic co-inhibition of Wnt and PI3K signalling improves survival and reduces tumour growth in cancer bearing mice. **A**. RNA sequencing data of human ICC demonstrating a positive correlation between the activity of canonical Wnt signalling and Akt signalling. **B**. Schematic representation of the KPPTom cholangiocarcinoma model where Cre^ERT^ expression in Keratin-19-positive cholangiocytes results in the inactivation of *Trp53* and *Pten*, whilst labelling transformed cells with tdTomato. **C**. immunohistochemistry showing that lineage traced ICC (tdTomato, upper panel) demonstrates activation of canonical Wnt signalling (by staining for dephosphorylated (active) β-catenin) and PI3K activity (through pAKT^Ser647^ positivity). Red arrows denote positive cells. Scale bar = 100 μm **D**. Immunohistochemical staining for tdTomato positive cancer cells in vehicle treated animals compared to those treated with a combination of LGK974 and Pictilisib. Scale bar = 100 μm. **E**. Number of tdTomato positive cells in KPPTom animals given vehicle or LGK974 and Pictilisib in combination. **F**. Proportion of KPPTom animals containing macroscopic tumours in KPPTom animals treated with vehicle vs combination treatment.

To test this, we deleted *Trp53*, which is the most common mutation in our patient cohort, mutated in 19.13% of cases (**Figure 1C**) and *Pten* specifically in biliary epithelial cells using a *Keratin19-Cre*^*ERT*^ knock-in mouse line. While *Pten* mutations were not found in our computational analysis of ICC, loss of *Pten* recapitulates the effects of PI3KCA mutations(46) (which is mutated in 5.4% of patients in our cohort, **Figure 1C**) resulting in the accumulation of PIP_3_ and AKT activation. Concurrently, with *Trp53* and *Pten* deletion, recombined cells were labelled with tdTomato (from here on in this line is known as the KPPTom line). When KPPTom mice are challenged with Thioacetamide, a hepatotoxin that has been used to induce ICC in other models but is not itself mutagenic(47,48), moderate to well-differentiated cholangiocellular neoplasms form around the portal tracts of mice within 8 weeks (Schematic in **Figure 4B** and **4C**). Tumours from the KPPTom mice are also positive for dephosphorylated (active) β-catenin, which localises to the cytoplasm and nucleus of cancer cells, as well as phosphorylated AKT^Ser647^ (**Figure 4D**). As KPPTom mice show both Wnt and PI3K/AKT activity, we treated them with LGK974 and Pictilisib to determine whether a model of well-differentiated ICC is susceptible to co-inhibition of these pathways. Following treatment, the number of tdTomato-positive cancer cells was significantly reduced (by 68.3%) when compared to control vehicle treated animals (**Figure 4E** and **4F**). Indeed when treated with LGK974 and Pictilisib, only 33% of KPPTom mice developed ICC, whereas in the vehicle treated cohort, 80% of KPPTom mice contained cancerous lesions (**Figure 4G**).

## Discussion

Intrahepatic cholangiocarcinoma (ICC) is highly complex at the genetic(28) and cellular levels(49). While a number of candidate mutations have been identified that can be pharmacologically targeted(19,50), these have not led to a broadly applicable treatment. Moreover, the presence of these mutations does not necessarily predict therapeutic responsiveness in patients with ICC and only a subset of these patients respond to targeted therapy(20). Consequently, there is a clinical necessity to identify the mechanisms that ICC uses to grow and define whether these processes can be used in patient stratification and targeted treatment. In order to define a targeted therapeutic approach that can be used in treating a specific cohort of ICC patients, we need to understand whether the genetic complexity found in ICC ultimately translates to phenotypic diversity or whether ICC relies on a limited number of signalling cascades to grow in spite of this genetic complexity.

The identification of causative mutations in ICC is fraught with complications. Historically these studies relied heavily on identification of genes with recurrent consensus mutations in patients. This approach has been severely limited by the relatively small number of ICC samples available(13,14,24,51). Increasing sample size by pooling data from across published cohorts and combining this with a driver prediction pipeline that puts less weight on mutation recurrence within a population, but rather concentrates on the patterns and predicted effects of mutations(16) enabled us to identify an expanded set of candidate mutations in ICC that have the potential to act as oncogenes.

Whilst able to predict novel candidate oncogenic mutations, the approach described here is not able to infer which mutations act in epistasis to promote tumour formation. To overcome this limitation we developed an *in vivo*, highly multiplexed CRISPR-SpCas9 screening approach. Rather than using whole genome screening as other studies have(52–54), we biased our libraries to ensure that they targeted genes mutated in ICC patients. Using this strategy has enabled us to define a range of putative tumour suppressors that cooperate with gain-of-function *Ras*-mutations to overcome RAS-induced senescence and initiate tumour formation. We validated that one of these, Neurofibromin-2 (*Nf2*, or Merlin) can act as an important factor in the initiation of RAS-driven ICC and we show increased penetrance and aggression of RAS-driven ICC when *Nf2*-loss is combined with *Trp53*-loss.

The phenotypes of these *Nf2*-mutant tumours are highly sarcomatoid, the cancer cells have a more spindle-like morphology and are dispersed throughout the liver. Perhaps this is unsurprising given the role of NF2 in maintaining contact inhibition and restricting cell migration in the presence of Ras mutations, as NF2 has been implicated in the fine tuning of Ras signalling in Schwann cells(55). However, loss of *Nf2* has not been shown to be sufficient to cooperate with Ras and drive tumour formation independent of other accessory mutations and while loss of *Nf2* has been previously shown to promote the formation of mixed HCC and ICC when it is deleted from the liver during development(56,57), it was unclear whether *de novo* somatic *Nf2* mutations interact with other ICC-relevant mutations *in vivo*. In addition to Hippo signalling(58), NF2 has also been implicated in the regulation of a number of other signalling pathways(34,59,60). We found that Wnt and PI3K signalling are recurrently de-regulated following *Nf2*-loss in our mouse model, leading us to identify co-inhibition of Wnt and PI3K as a potent therapeutic in reducing ICC growth and improving survival in mouse models of ICC which are both dependent on or independent of *Nf2* status. Both Wnt (porcupine) inhibitors and PI3K inhibitors are currently in clinical trials for other solid malignancies and our data supports recent findings in pancreatic cancer(61), that treating patients with Wnt and PI3K inhibitors provides an attractive therapeutic route to reduce tumour burden, particularly now the known side-effects of Wnt-inhibition can be managed(62). Our data demonstrates the importance of understanding the function of rare mutations in ICC and shows that these low-frequency mutations not only influence the outcome of more common driver mutations, but also can lead us to define applicable therapeutic strategies that can be used to develop personalised approaches which could be applied clinically to stratify patients and treat ICC of divergent genotypes based on the signalling pathways that are deregulated in these cancers.

## Materials and Methods

### Identification and processing of genomic data from patient datasets

#### Alignment and pre-processing of publicly available data

Exome-seq FASTQ files from Chan-on et al. (14). were downloaded from the European Nucleotide Archive with accession PRJEB4445. Exome-seq FASTQ files from Sia et al. (23) were downloaded from the Gene Expression Omnibus Database with accession GSE63420. TCGA BAM files were downloaded from the Genomic Data Commons after receiving access to individual patient BAM files.

For the TCGA, Chan-On, and Sia datasets, mutations were called as follows. Input FASTQ files were aligned to the Hg19 reference genome using Burrows-Wheeler Aligner version 0.7.15(63). PCR and optical duplicates were marked using PicardMarkDuplicates version 2.5.0, base quality score recalibration was carried out using BaseRecalibrator and local indel realignment was performed with IndelRealigner (both from the Genome Analysis Toolkit(64) version 3.6 (GATK3)). Ensemble variant calling was performed with Mutect (1.1.5)(65), Mutect2 (2.1)(66), Freebayes (1.0.2.29)(67), Vardict-java (1.4.6)(68), and Varscan (2.4.2)(69) and all passing somatic variants which were identified by 2 or more algorithms were taken forward. The above analysis was batched and implemented within the Blue Collar Bioinformatics pipeline (bcb io-nextgen 1.0.0a0-af4730e). TCGA data were input as reads which were aligned to the Hg38 reference so, following mutation calling, variants were re-mapped using Picard LiftoverVCF (2.5.0) using the Hg38toHg19 chain file provided by UCSC. Variant annotation was performed with the Ensembl Variant Effect Predictor (v88) using the ‘--pick’, ‘--tab’, and ‘--symbol’ options.

#### Identification of candidate pathogenic variants using IntOgen

IntOgen(16) was run on each published/TCGA cohort individually, and then all combined. Significance thresholds for OncoDriveFM and OncoDriveCLUST were q=0.05, and q=0.1 for MutSigCV as per the default recommendations. The minimum frequency of occurrence for a gene to be considered for analysis was n=2 for OncoDriveFM and n=5 for OncoDriveCLUST. Following candidate identification with IntOgen, functional Interaction Inference Cytoscape(25) (3.7.1) with the ReactomeFIVis(26) app was used to build a network of known and inferred functional interactions between the set of predicted drivers so as to explore the relationship between them in a cellular context. This network was clustered into modules by connection density and each module was annotated with pathway enrichments.

### Design and preparation of sgRNA plasmids for in vivo editing

#### Generation of pooled sgRNA screening library

Oligonucleotides encoding sgRNAs targeting the set of predicted drivers were designed using spacer sequences from the mouse GeCKo V2 library with the Python script supplied in the corresponding paper (**Supplementary Materials Table 1**)(70). Library-specific PCR retrieval arms were derived from the protocol for parallel oligonucleotide retrieval published by the Luo lab(71) and our schematic for library preparation is described in **Supplementary Materials Table 1**. Complete sgRNA oligos for all target genes and control sequences were custom synthesised by Twist Biosciences. Library oligos were amplified from the custom oligo pool using PCR for 22 cycles and products were cleaned-up using the QIAquick Nucleotide Removal Kit. Purified amplicons were then digested with Esp3I and phosphorylated. Digested inserts were purified from the reaction by isopropanol precipitation and ligated into the SB-CRISPR plasmid backbone overnight at 16°C with T4 ligase. Pools were transformed into Stbl3 to ensure unbiased and complete library representation. Transformants were grown overnight at 37°C. Plasmids were isolated using a Maxiprep Kit as per the manufacturer’s instructions. Resultant libraries underwent amplicon sequencing to determine the representation and evenness in sgRNA sequences. sgRNA inserts were PCR amplified with primers targeting the sequence before and after the insert site. This step also served to add MiSeq (Illumina) flow-cell adaptors, variable length random sequences to increase sequencing library complexity, and index sequences for multiplexed sequencing.

#### Generation of single gRNAs

Single gRNAs (**Supplementary Materials Table 2**) were cloned into SB-CRISPR plasmids kindly provided by Professor Dr Roland Rad (LMU Munich). Oligos encoding sgRNAs against *Trp53* and *Nf2* or non-targeting control gRNA (‘0007’) were cloned into the backbones as follows. SB-CRIPSR was digested with Esp3I or BbsI to produce staggered ends, with simultaneous dephosphorylation using FastAP Thermosensitive Alkaline Phosphatase to prevent backbone re-closure during ligation. Plasmid digestion products were size separated by gel electrophoresis in 2% agarose gel following which it was extracted and purified. sgRNA oligos were annealed and simultaneously phosphorylated with T4 Polynucleotide Kinase. Ligation of plasmid backbone and insert was carried out at 16°C overnight with T4 and then transformed into Stbl3 cells as per the manufacturer’s instructions and grown at 37°C overnight. Sanger sequencing of colonies identified those with correctly ligated plasmids which were subsequently picked, cultured and extracted as maxi-prep’s as per the manufacturer’s instructions. Plasmid concentrations were quantified using a Nanodrop Spectrophotometer.

#### Animal work

All animal work was performed under the UK Home Office project license held by Dr Luke Boulter (PFD31D3D4). Animals were maintained in colonies in 12h light-dark cycles and were allowed access to food and water *ad libitum*.

#### Hydrodynamic tail vein injection

Female, FVB/N mice were purchased from Charles River, UK and were used at 4-6 weeks of age. For the hydrodynamic tail vein injection, animals were injected with a physiological saline solution (10% w/v) containing plasmids into the lateral tail vein in <7 seconds to achieve a hydrodynamic transfection of the liver. The typical injection contained 6μg of PGK-SB13, 20ug of CAG-Kras^G12D^ or CAG-Nras^G12V^ and 20μg of SB-CRISPR gRNA plasmid or shRNA^Trp53-GFP^ plasmid. In models that relied on a combination of gRNA plasmids, plasmids were dissolved to a maximum concentration of 20μg and were mixed such that they were balanced pools of each gRNA. For screening studies, the gRNA library was injected at 20ug and we determined gRNA representation by Sanger sequencing prior to injection and plotted the GINI index for each library **(Supplementary Figure 4**).

#### *Keratin-19-CreER*^*T*^;*Pten*^flox/flox^;*Trp53*^flox/flox^;R26R^LSLtdTomato^ mice (KPPTom)

*Keratin-19-CreER*^*T*^ mice (Jax: 026925) were crossed with animals containing floxed alleles of *Pten* (Jax: 006440) or *Trp53* (Jax: 008462) and a silenced tdTomato reporter targeted to the Rosa26 locus (Jax: 007908) . All animals in this study are heterozygous for Keratin-19CreER^T^, homozygous for Trp53^flox^ and Pten^flox^ alleles and homozygous for R26R^LSLtdTomato^. Mice received three doses of 4mg of tamoxifen by oral gavage and followed by 400mg Thioacetamide in their drinking water. Animals developed well-differentiated bile duct adenocarcinoma within 8 weeks.

#### Therapeutic dosing of animal models

Animals baring KRAS^G12D^;*Trp53*^KO^;*Nf2*^KO^ tumours were dosed with either vehicle alone (10% DMSO, 40% PEG300, 5% Tween-80 and 45% saline), 5mg/Kg LGK974, 50mg/Kg Pictilisib or a combination of the two daily. These mice were dosed with these drug combinations starting 7 days following the hydrodynamic injection. KPPTom animals were given tamoxifen and Thioacetamide (as detailed above) and at four weeks were given either vehicle alone (10% DMSO, 40% PEG300, 5% Tween-80 and 45% saline) or a combination of 5mg/Kg LGK974 and 50mg/Kg Pictilisib for 4 weeks.

#### Isolation of RNA and DNA

Both DNA and RNA extraction used 50-100mg of snap frozen tissue. DNA was extracted from tissue using the DNeasy Blood and Tissue Kit (QIAGEN) as per the manufacturer’s instructions. RNA was extracted by placing the tissue into TRIzol RNA Isolation Reagent (Invitrogen), which was then homogenized with a steel bead in a Qiagen TissueLyser LT (QIAGEN). RNA was precipitated with chloroform and cleaned up using the RNeasy Mini Kit (QIAGEN) as per the manufacturer’s instructions. For downstream sequencing applications DNA and RNA quality (RIN score) was quantified using the Agilent 2100 Bioanalyzer with either the DNA 1000 chip, or RNA 6000 chip. A minimum RIN threshold of 8 was used for RNA-seq.

#### RNA Sequencing

Libraries were prepared from total-RNA samples using the NEBNext Ultra 2 Directional RNA library prep kit for Illumina with the NEBNext rRNA Depletion kit according to the provided protocol. 500ng of total-RNA was added to the ribosomal RNA (rRNA) depletion reaction using the NEBNext rRNA depletion kit (Human/mouse/rat). This step used specific probes that bind to the rRNA in order to cleave it. rRNA-depleted RNA was then DNase treated and purified using Agencourt RNAClean XP beads (Beckman Coulter Inc). RNA was then fragmented using random primers before undergoing first strand and second strand synthesis to create cDNA. cDNA was end-repaired before ligation of sequencing adapters, and adapter-ligated libraries were enriched by 10 cycles of PCR using NEBNext Multiplex oligos for Illumina. Final libraries had an average peak size of 260bp. Library QC: Libraries were quantified by fluorometry using the Qubit dsDNA HS assay and assessed for quality and fragment size using the Agilent Bioanalyser with the DNA HS Kit (#5067-4626). Fragment size and quantity measurements were used to calculate molarity for each library. Sequencing: Sequencing was performed using the NextSeq 500/550 High-Output v2.5 (150 cycle) Kit on the NextSeq 550 platform (Illumina Inc). Libraries were combined in an equimolar pool based on Qubit and Bioanalyser assay results and run across a single High Output v2.5 Flow Cell.

#### RNA sequencing data processing and analysis

The primary RNA-Seq processing, quality control to transcript-level quantitation, was carried out using nf-core/rnaseq v1.4.3dev (https://github.com/ameynert/rnaseq)(72). Reads were mapped to the mouse FVB_NJ_v1 decoy-aware transcriptome using the salmon aligner (1.1.0). RNA-Seq analysis was performed in R (4.0.2), Reads were summarized to gene-level and differential expression analysis was performed using the bioconductor packages tximport (1.16.1) and DESeq2 (1.28.1). A pre-filtering was applied to keep only genes that have at least 10 reads in a group and 15 reads in total. The Wald test was used for hypothesis testing for pairwise group analysis. A shrunken log2 fold changes (LFC) was also computed for each comparison using the adaptive shrinkage estimator from the ‘ashr’ package.

#### DNA exome sequencing

Libraries were prepared from each genomic DNA (gDNA) sample using the SureSelect XT Target Enrichment System for Illumina Paired-end Multiplexed Sequencing Library kit (according to the provided protocol (Version C2, Dec 2018). 200ng of each DNA sample was sheared using the Covaris E220e (Covaris) to achieve target DNA fragment sizes of between 150 and 200bp. DNA fragments were then end-repaired and purified using AMpure XP beads (Beckman Coulter). After end-repair a single ‘A’ nucleotide was added to the 3’ ends of the blunt fragments to prevent them from ligating to each other during the subsequent adapter ligation reaction. Adapter-ligated libraries were purified using AMPure XP beads to remove leftover adapters and adapter-dimers prior to amplification for 10 cycles of PCR. After a final AMPure XP cleanup, gDNA libraries were quantified with the Qubit dsDNA HS assay and quality was assessed on the Agilent Bioanalyser (Agilent Technologies Inc) with the DNA High Sensitivity kit (#5067-4626). 750ng of each prepared gDNA library was hybridised overnight to probes designed to cover the mouse exome using the SureSelect XT Mouse All Exon Capture Library (Agilent Technologies). Dynabeads MyOne Streptavidin T1 magnetic beads (ThermoFisher Inc) were used to capture the hybridised DNA-probes, and a series of washes were undertaken to remove non-hybridized DNA. Captured libraries were then amplified for 12 cycles of PCR with unique indexing primers to allow multiplexed sequencing before a final cleanup with AMPure XP beads. Library QC: Libraries were quantified by Qubit using the dsDNA HS assay and assessed for quality using the Agilent Bioanalyser DNA HS Kit. Molarity for sequencing was calculated using the Qubit results and library fragment size information from the Bioanalyser traces. Sequencing: Sequencing was performed using the NextSeq 500/550 High-Output v2.5 (150 cycle) Kit on the NextSeq 550 platform (Illumina Inc). Libraries were combined in a single equimolar pool and run on a High-Output v2.5 Flow Cell.

#### CRISPR/Cas9-editing validation and Structural Variant calling

DNA sequences were extracted from FASTQ files from exome-sequencing of tumours arising from the RAS^G12^-library screens and were aligned to the FVB mouse reference genome; subsequently, indels within 50bp upstream or downstream of sgRNA target sites were called. To determine if indels were likely due to SpCas9 activity or were spurious mutations, the interval of each indel was observed on the FVB genome using Integrative Genomics Viewer(73), and overlain with sgRNA library binding sites. Indels with start or end sites out-with sgRNA targets were removed, although this was only the case for 5 indels across all samples. To determine if editing had induced structural rearrangements, structural variants (SVs) were called using Delly2 (v0.7.9) (74)as follows: all reads which had a minimum mapping quality of 20 were used to call germline and somatic SVs, SVs with a PASS flag for somatic status, had a minimum variant allele fraction of 0.15, had zero read support in the normal, and had precisely mapped breakpoints were then visually analysed on IGV to determine if they overlapped with sgRNA target sites.

#### Data curation and deposition

All RNA and Exome sequencing data pertaining to this manuscript is deposited on the NCBI Gene Expression Omnibus (GEO) as accession number GSEXXXXXX.

#### Histology and Immunohistochemistry

Livers were perfused with phosphate buffered saline and dissected into 10% neutral buffered formalin. Fixed tissue was processed in wax blocks and sectioned 4 μm thick. Sections form immunostaining were dewaxed in xylene and rehydrated. Following antigen retrieval (**Supplementary Materials Table 3**) samples were incubated with 3% hydrogen peroxide and endogenous Avidin and Biotin were blocked. Non-specific protein binding was blocked using a pan-species protein block. Primary antibodies were diluted in antibody diluent and incubated overnight. Primary antibodies were detected using either a species specific biotinylated secondary antibody and HRP-DAB detection system or directly conjugated fluorescent secondary antibodies (**Supplementary Materials Table 3**). Slides were either counterstained with Harris haematoxylin in the case of DAB staining or mounting media containing DAPI for fluorescent imaging. Histological assessment was undertaken by a consultant liver histopathologist working at the national liver transplant centre (TJK) with experience in the comparative pathology of animal models of primary liver cancer.

#### Quantification of tumour burden

Histological sections containing tumours were scanned using a Nanozoomer slide scanner with a 40X objective lens. Files were then imported into QuPath (https://qupath.github.io) and tumour tissue was manually annotated. The total area of the liver was also captured in this process. Tumour Burden represents the area of tissue occupied by tumour and number is the number of discrete tumours in the tissue.

#### Reverse phase protein arrays

Snap frozen, dissected tumour tissue was provided to the Human Tumour Profiling Unit (HTPU) at the Cancer Research UK Edinburgh Centre. The target proteins analysed by RPPA are listed in **Supplementary Materials Table 4**. Reverse Phase Protein Array (RPPA) analysis was carried out using established protocols for nitrocellulose-based arrays(75). Briefly, cell lysates were prepared in 1% Triton X-100, 50 mM HEPES (pH 7.4), 150 mM sodium chloride, 1.5 mM magnesium chloride, 1 mM EGTA, 100 mM sodium fluoride, 10 mM sodium pyrophosphate, 1 mM sodium vanadate, 10% glycerol, supplemented with complete ULTRA protease inhibitor and PhosSTOPTM phosphatase inhibitor cocktails (Sigma Aldrich). After clearing by centrifugation at 13,300□rpm for 10□min at 4°C the protein concentration was determined using Coomassie Plus Protein Assay (ThermoFisher Scientific). Protein concentrations were adjusted to a final concentration of 2mg/ml followed by denaturation upon addition of 4X sample buffer containing 10% Beta-mercaptoethanol and heating to 95oC for 5 minutes. A 4-step dilution series of each sample was prepared in PBS with 10% glycerol giving final concentrations of 1.5, 0.75, 0.375 and 0.1875 mg/ml. Samples were printed onto nitrocellulose-coated slides (Grace Bio-Labs) across multiple sub-array areas under conditions of constant 70% humidity using an Aushon 2470 array platform (Quanterix). Printed slides were blocked using SuperBlock (TBS) blocking buffer (Thermo Fisher Scientific) and each sub-array was separately incubated with validated primary antibodies (all diluted 1:250 in SuperBlock). Bound antibodies were detected by incubation with anti-rabbit or anti-mouse DyLight 800-conjugated secondary antibodies (New England BioLabs). Slide images were acquired using an InnoScan 710-IR scanner (Innopsys) with laser power and gain settings optimised for highest readout without saturation of the fluorescence signal. The relative fluorescence intensity of each array feature was quantified using Mapix software (Innopsys).

The linear fit of the dilution series of each sample was verified for each primary antibody and the median relative fluorescence intensity from the dilution series was calculated to represent relative abundance of total protein and post-translational epitopes across the sample set. Finally, signal intensities for each sample were normalized to total protein loading on the array slides by using the signal readout from a fast-green (total protein) stained array.

#### Statistical Analysis

All experimental groups were analysed for normality using a D’Agostino– Pearson Omnibus test. Groups that were normally distributed were compared with either a two-tailed Student’s t test (for analysis of two groups) or using one-way ANOVA to compare multiple groups, with a post hoc correction for multiple testing. Non-parametric data were analysed using a Wilcoxon– Mann–Whitney U test when comparing two groups or a Kruskall–Wallis test when comparing multiple non-parametric data. Throughout, P<□0.05 was considered significant. Data are represented as mean with S.E.M. for parametric data or median with S.D. for non-parametric data.

## Acknowledgements

NTY and MLW designed and performed experiments, analysed the data and prepared the manuscript. AMM analysed and prepared data for publication. KG, EJJ, SHW, PAT and DHW performed experiments and analysed data. RVG, SJW, JCA provided tissue and generated reagents used in this study. TJK provided clinical pathological support throughout the project, DS, PM, MST provided scientific direction and edited the manuscript. LB funded the project, designed and performed experiments, analysed data and wrote the manuscript. LB is funded by The Wellcome Trust (207793/Z/17/Z), AMMF (2016/108, 2017/115) and Cancer Research UK (C52499/A27948).

## Supplementary Material

Supplementary Figure 1 – Sequencing analysis pipeline

Supplementary Figure 2 – Cohort Demographics

Supplementary Figure 3 – Library generation and representation

Supplementary Figure 4 – Screened tumours are poorly differentiated adenocarcinomas with glandular features

Supplementary Figure 5 – Outcome of CRISPR-SpCas9 editing events

Supplementary Figure 6 – Analysis of CRISPR-induced edits per tumour in exome-sequenced cancer

Supplementary Figure 7 – Cancers containing Nf2-loss tend to segregate differently based on their transcriptomes

Supplementary Figure 8 – Clustering of RNAseq data and PANTHER analysis

Supplementary Figure 9 – The combined loss of *Nf2* and *Trp53* results in suppression of cancer cell apoptosis

Supplementary Figure 10 – Segregated survival data for animals treated with Wnt and PI3K inhibitors

Supplementary materials tables 1-4 – containing information about experimental design in this study

Supplementary table 1 – Identification and removal of hypermutated samples

Supplementary table 2 – Human data sources for driver prediction

Supplementary table 3 – Driver-Sample matrix

Supplementary table 4 – IntOGen results across sources

Supplementary table 5 – Percentage of cohort containing drivers

Supplementary table 6 – Gene ontology per module

Supplementary table 7 – Differentially expressed genes between groups

Supplementary table 8 – Correlation between AKT and Wnt signatures in human disease

