## Supplementary figures and images for "*In vivo* modelling of patient genetic heterogeneity identifies concurrent Wnt and PI3K activity as a potent driver of invasive cholangiocarcinoma growth"

### Supplementary Table 6

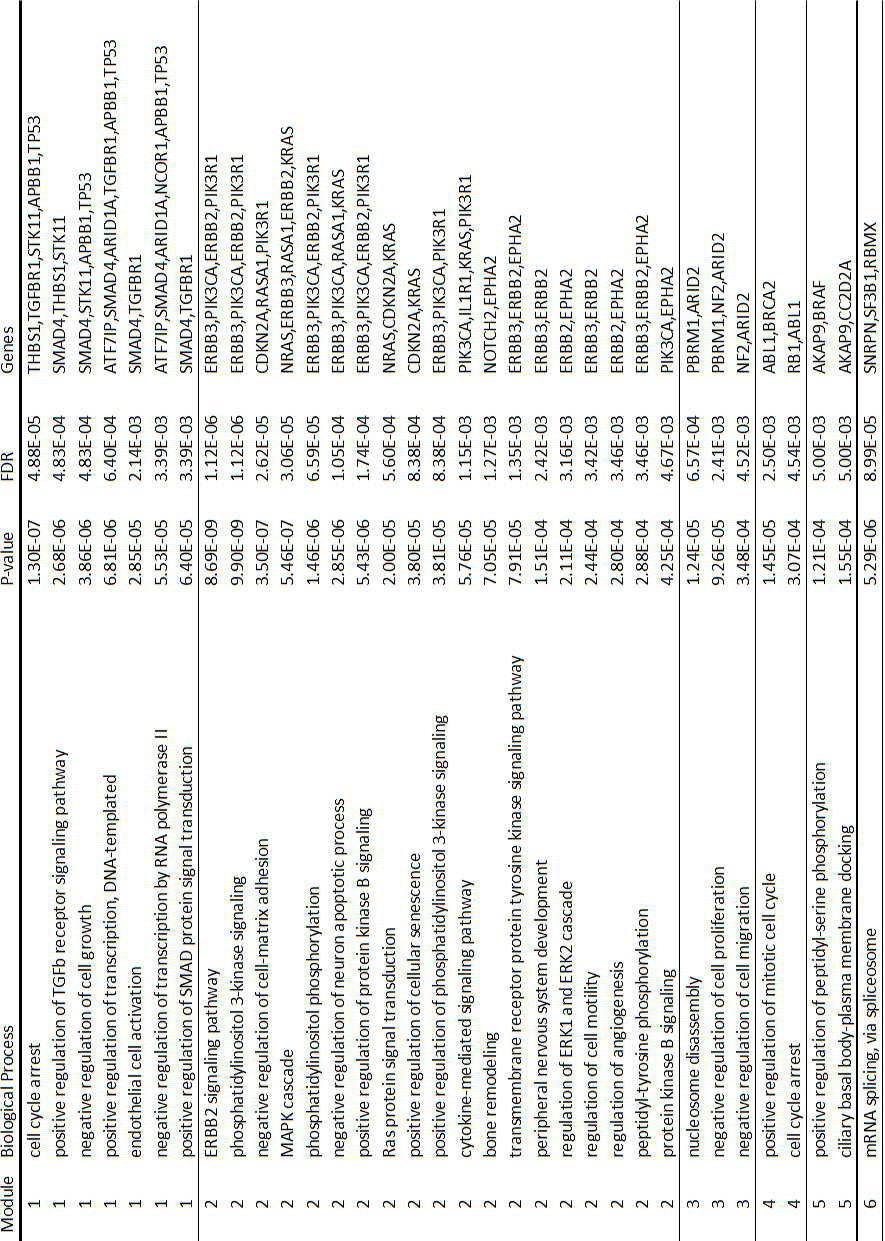
