## Supplementary Figures for "*In vivo* modelling of patient genetic heterogeneity identifies concurrent Wnt and PI3K activity as a potent driver of invasive cholangiocarcinoma growth"

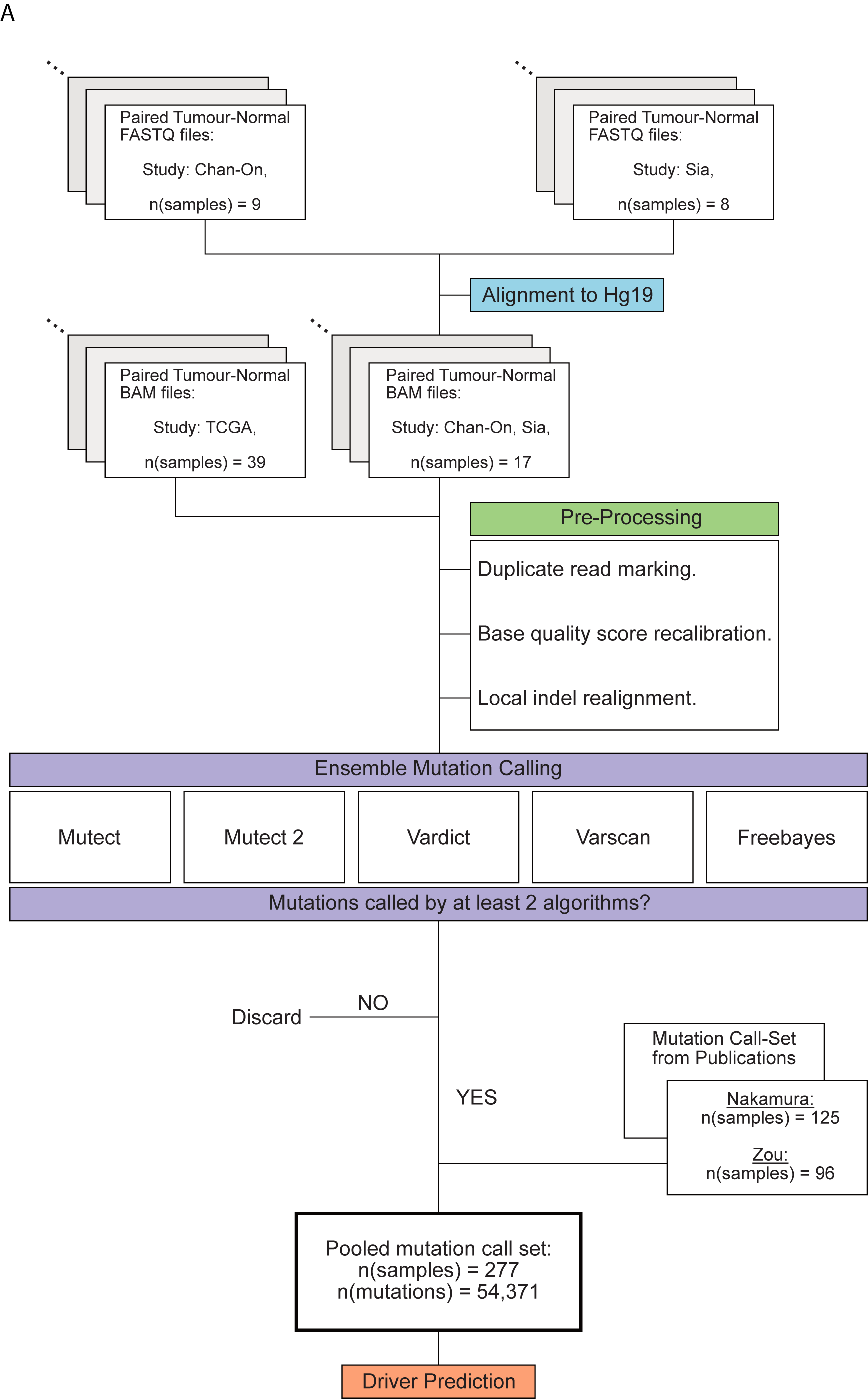
**Supplementary Figure 1 – Sequencing analysis pipeline:** **A**. Schematic representation of the data processing and data analysis pipeline applied to this study.

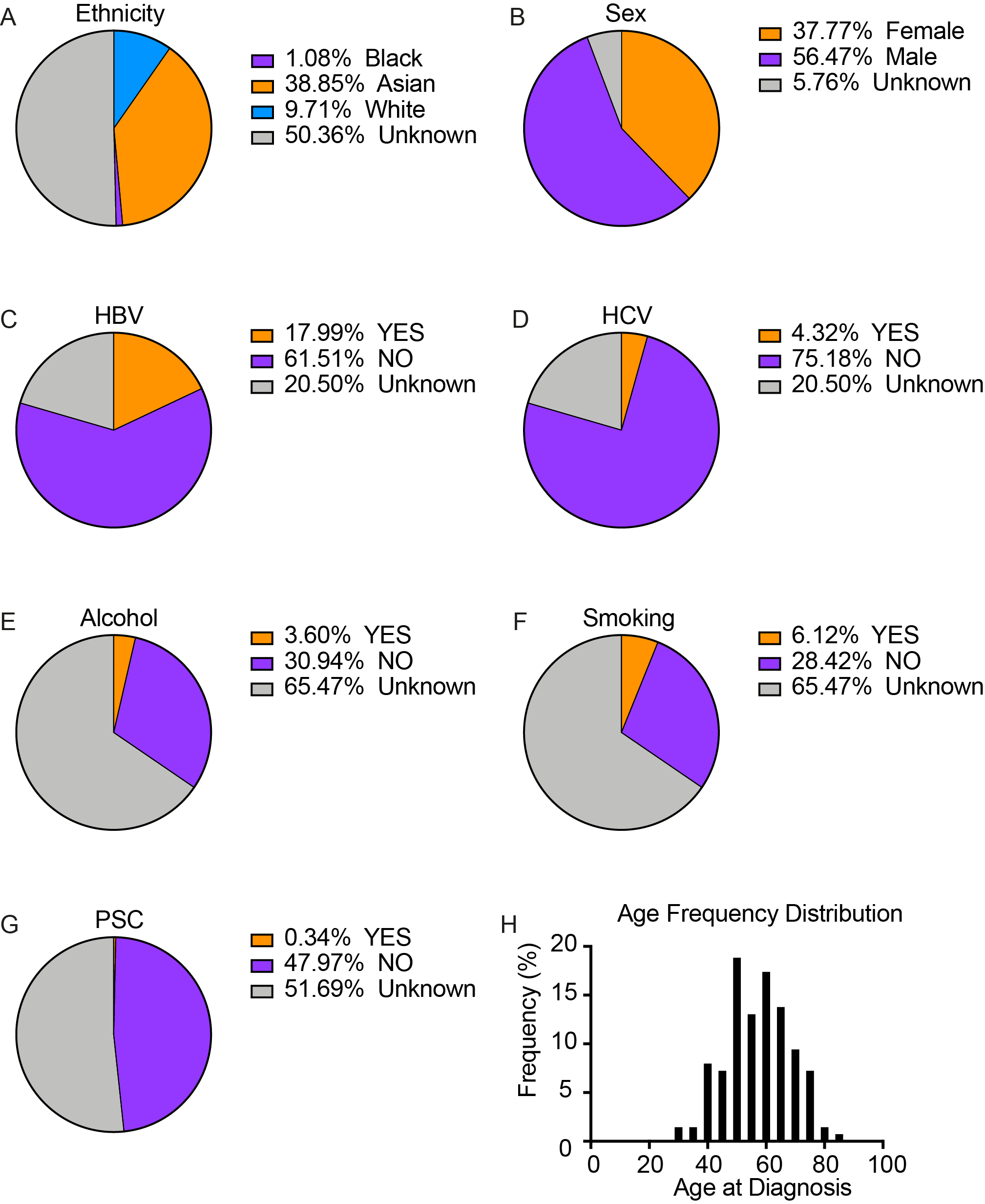
**Supplementary Figure 2 – Cohort Demographics:** **A.** Ethnic distribution of participants in this study (for those where the data is known). **B-G.** The proportion of each clinical attribute in the combined cohort. **H.** Distribution of ages of patients in the pooled cohort, the average is centred around 57 years.

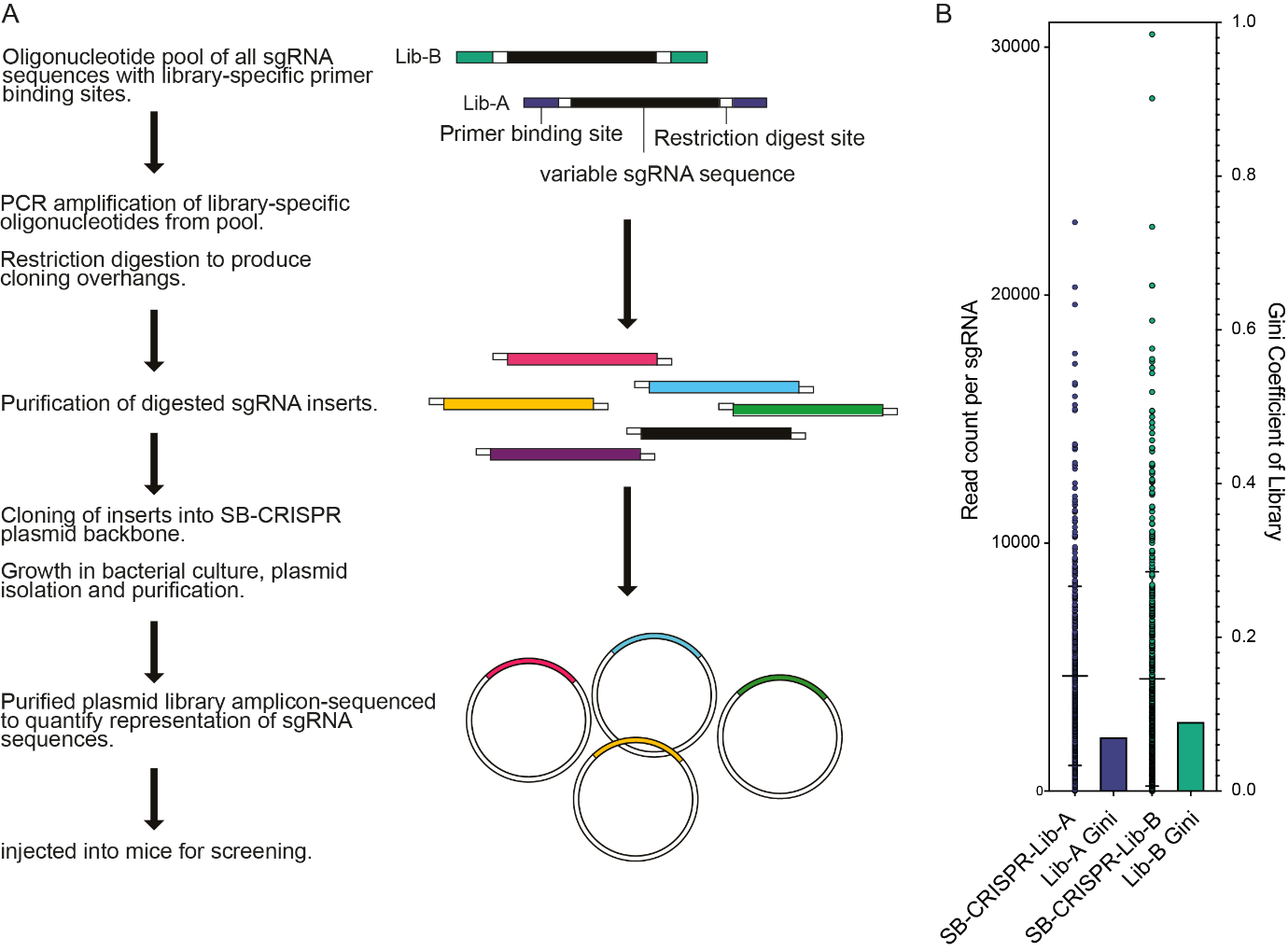

**Supplementary Figure 3 – Library generation and representation.** **A.** schematic of how the multiplexed ICC^Lib^ library is made; libraries are amplified from a pool and cloned into the SB-CRISPR backbone. **B.** MiSeq amplicon sequencing showed a low Gini index indicative of equal sgRNA sequence representation.

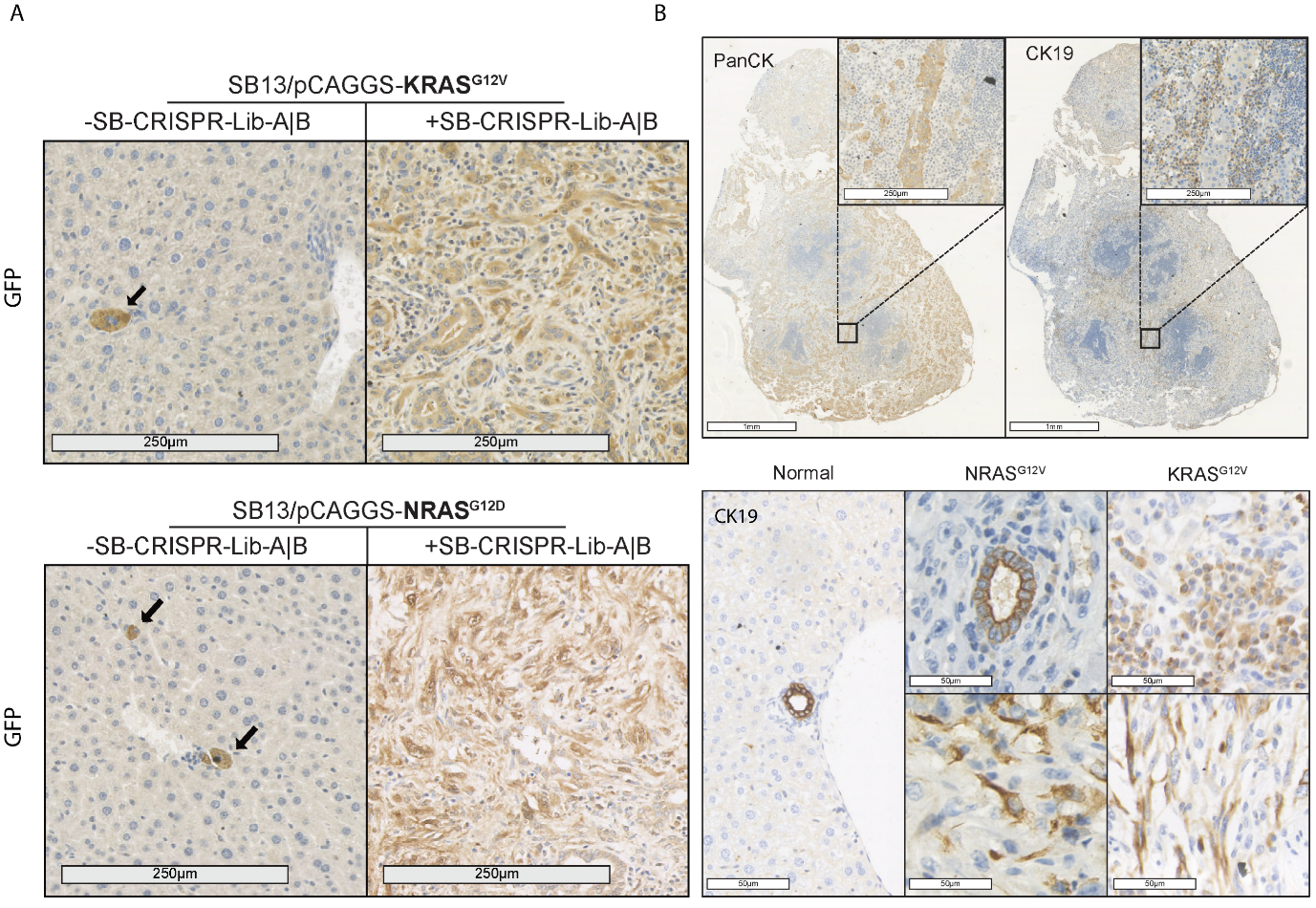

**Supplementary Figure 4 – Screened tumours are poorly differentiated adenocarcinomas with glandular features: A.** Immunohistochemistry for GFP (NRAS^G12V^ and KRAS^G12D^) expressing cells in mouse livers containing either a control vector (left panels) or SB-CRISPR Libraries (right panels). **B.** Immunohistochemistry of screened tumours for the epithelial marker pan-Cytokeratin (panCK), upper panels and the biliary lineage marker Cytokeratin-19 (CK19), lower panels in KRAS^G12D^ expressing tumours.

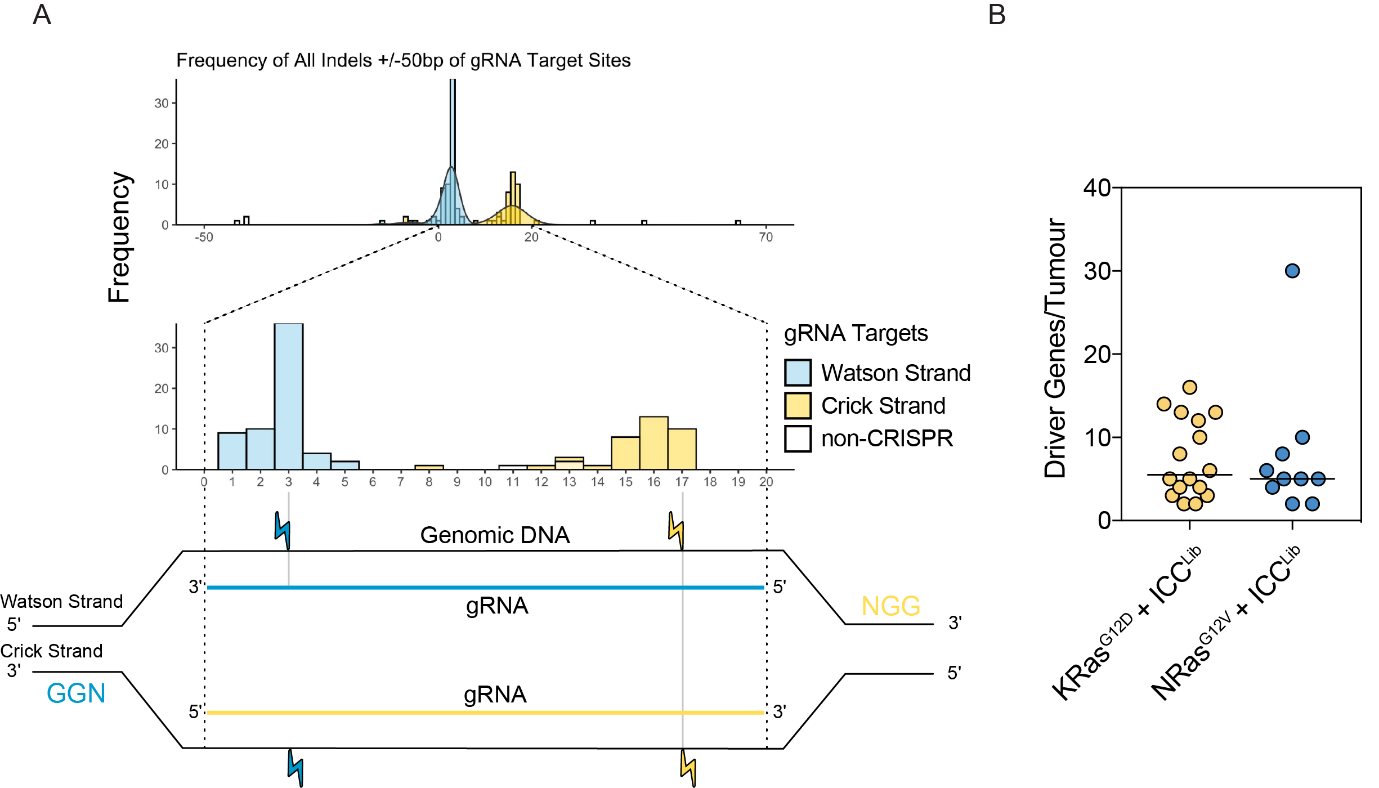

**Supplementary Figure 5 – Outcome of CRISPR-SpCas9 editing events:** **A.** indels within 50bp of sgRNA spacer target sites localise 2-4bp proximal to PAM sequences indicative of CRISPR/Cas9 mediated editing. **B.** The number of driver genes per tumourin in NRAS^G12V^ and KRAS^G12D^ oncogene screens when co injected with the ICC^Lib^.

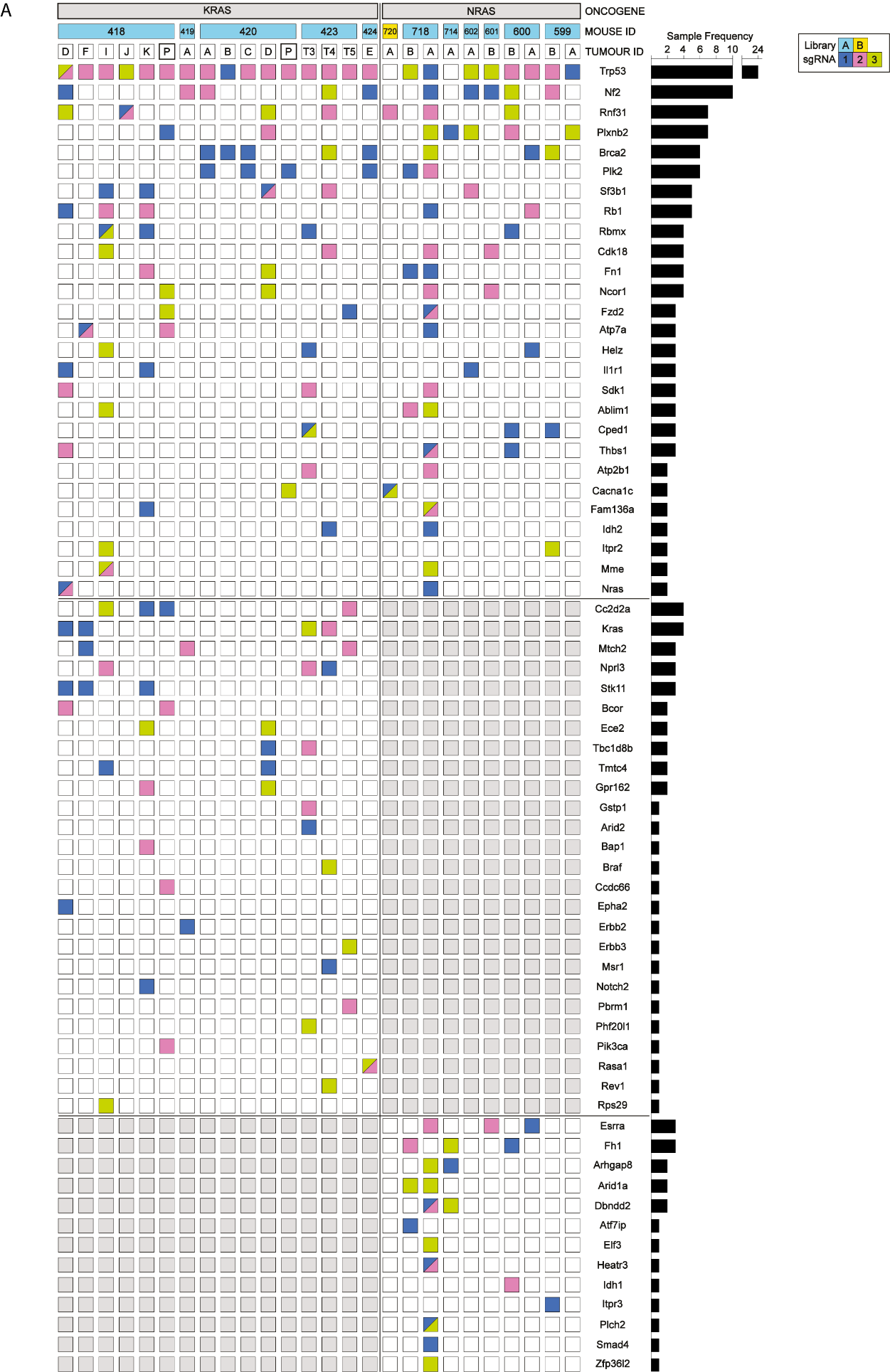

**Supplementary Figure 6 – Analysis of CRISPR-induced edits per tumour in exome-sequenced cancer: A.** Tabulated results showing which genes are mutated in each of 26 tumours generated through the co-injection of either oncogenic NRAS^G12V^ or KRAS^G12D^ along with gRNA libraries targeting mutant ICC genes.

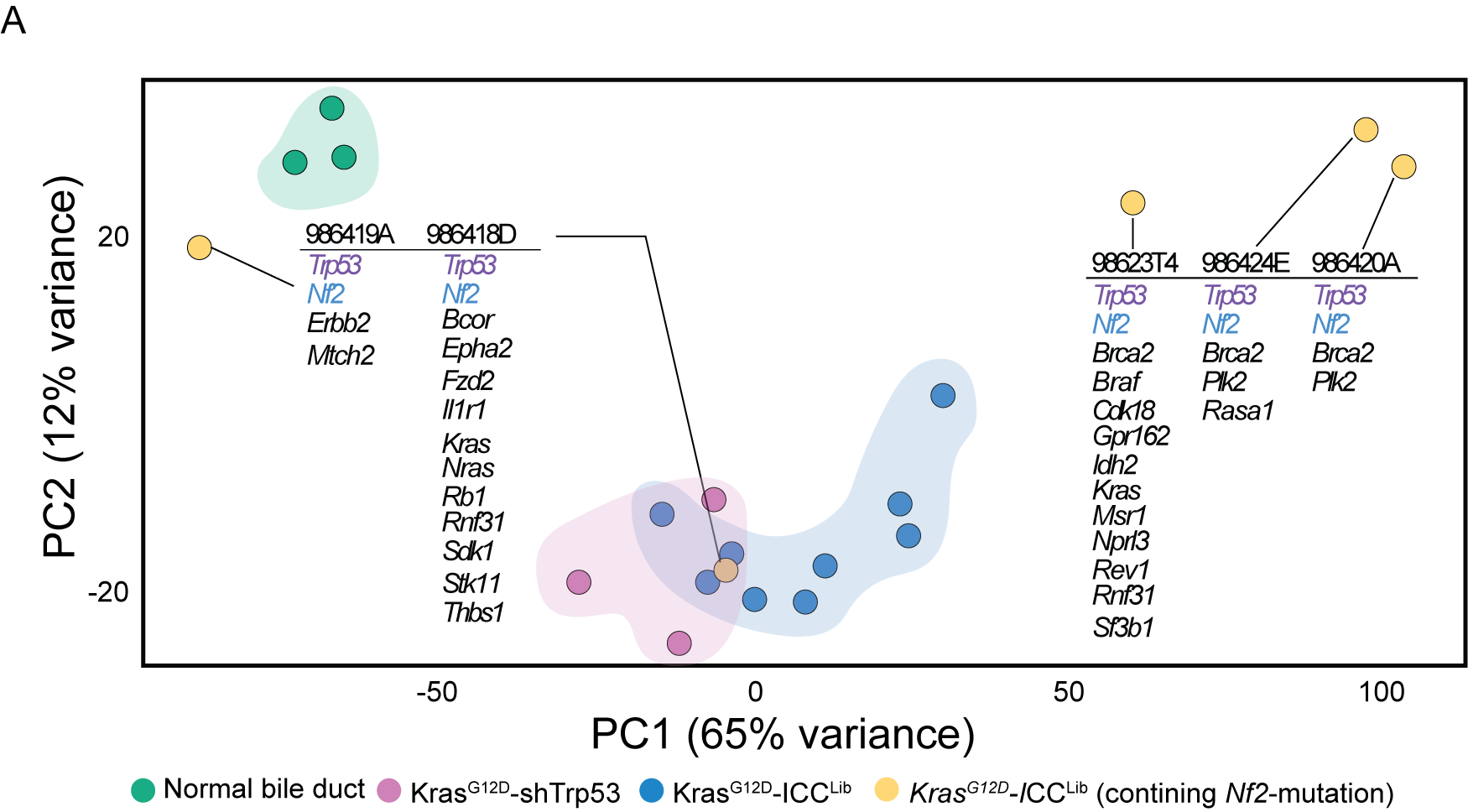

**Supplementary Figure 7 – Cancers containing *Nf2*-loss tend to segregate differently based on their transcriptomes.** A. Principal Component Analysis showing how samples group based on their transcriptomic signature. Normal bile ducts (green) cluster together. The majority of screen tumours (blue) cluster closely to tumours driven by KRAS^G12D^ and a shRNA targeting *Trp53* (magenta), except for those with *Nf2*-mutations, which form independent clusters (yellow).

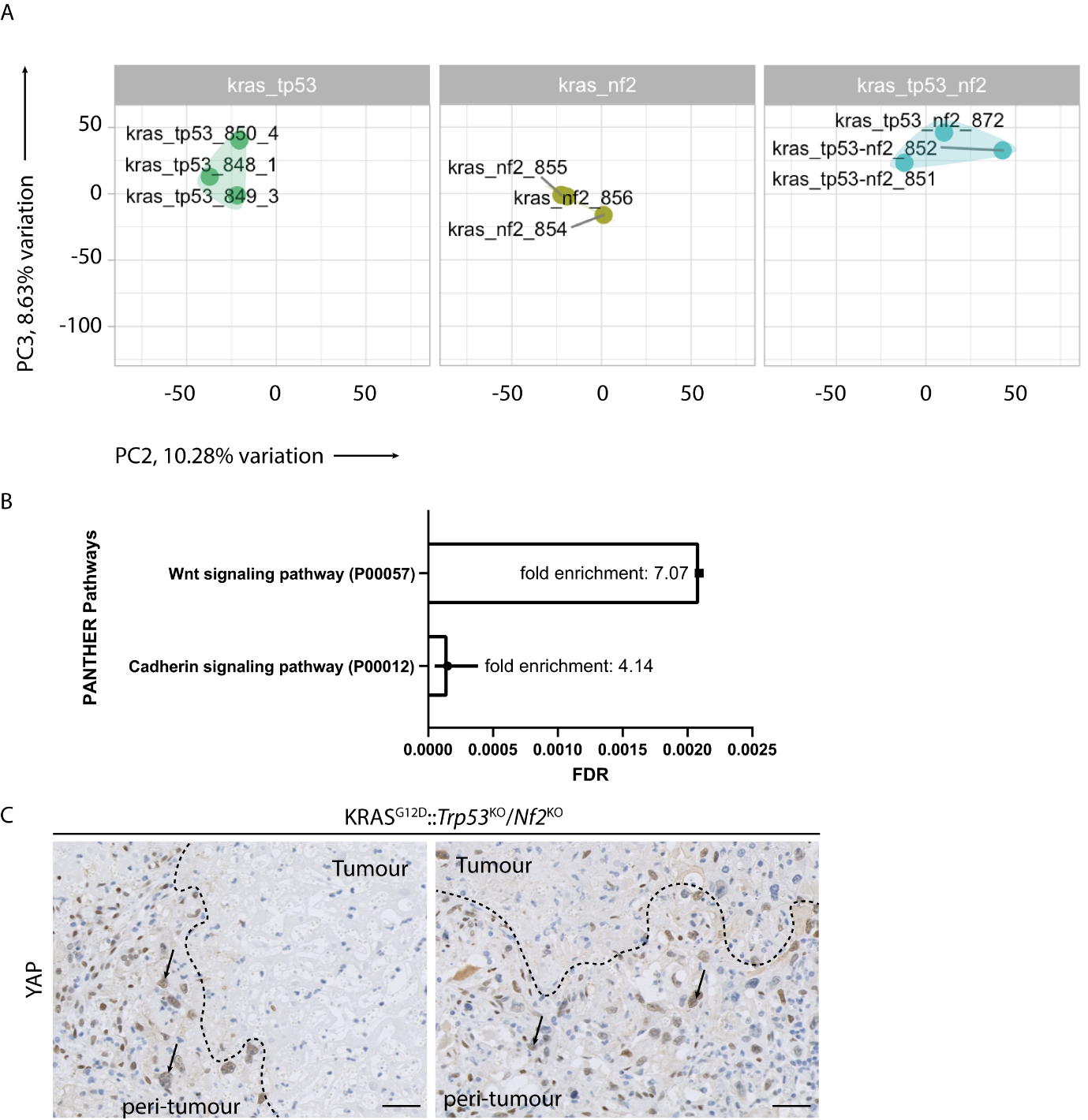

**Supplementary Figure 8 – Clustering of RNAseq data and PANTHER analysis:** **A.** Principal Component analysis of whole transcriptomes from KRAS^G12D^;*Trp53*^KO^, KRAS^G12D^;*Nf2*^KO^, KRAS^G12D^;*Trp53*^KO^;*Nf2*^KO^ tumours. **B.** Output of GO term analysis of significantly up and down regulated transcripts that are shared between groups when KRAS^G12D^;*Trp53*^KO^ is compared to KRAS^G12D^;*Trp53*^KO^/*Nf2*^KO ­^and when KRAS^G12D^;*Nf2*^KO^ is compared to KRAS^G12D^;*Trp53*^KO^;*Nf2*^KO^. **C.** Immunohistochemistry for dephosphorylated YAP in *Nf2*-deleted cancers. Black arrows denote positive nuclei, dotted line denotes the boundary between tumour and non-tumour tissue (labelled peri-tumour). Scale bar = 50 μm.

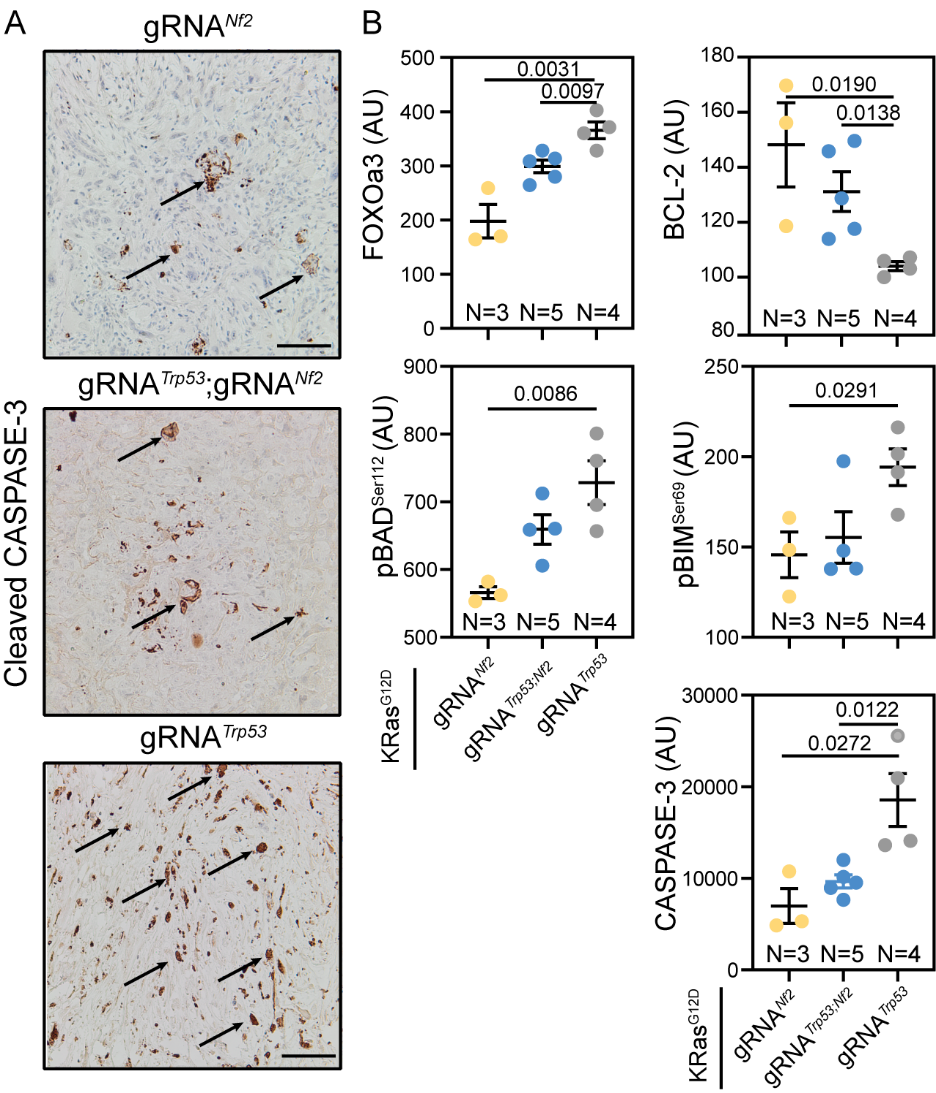

**Supplementary Figure 9 – The combined loss of *Nf2* and *Trp53* results in suppression of cancer cell apoptosis.** **A.** Immunohistochemistry staining for the apoptosis marker cleaved Caspase-3 on KRAS^G12D^;*Trp53*^KO^, KRAS^G12D^;*Nf2*^KO^, KRAS^G12D^;*Trp53*^KO^;*Nf2*^KO^ tissues. (Scale bar = 200 μm) **B.** RPPA analysis of apoptotic proteins using KRAS^G12D^;*Trp53*^KO^, KRAS^G12D^;*Nf2*^KO^, KRAS^G12D^;*Trp53*^KO^;*Nf2*^KO^ tumours as input material.

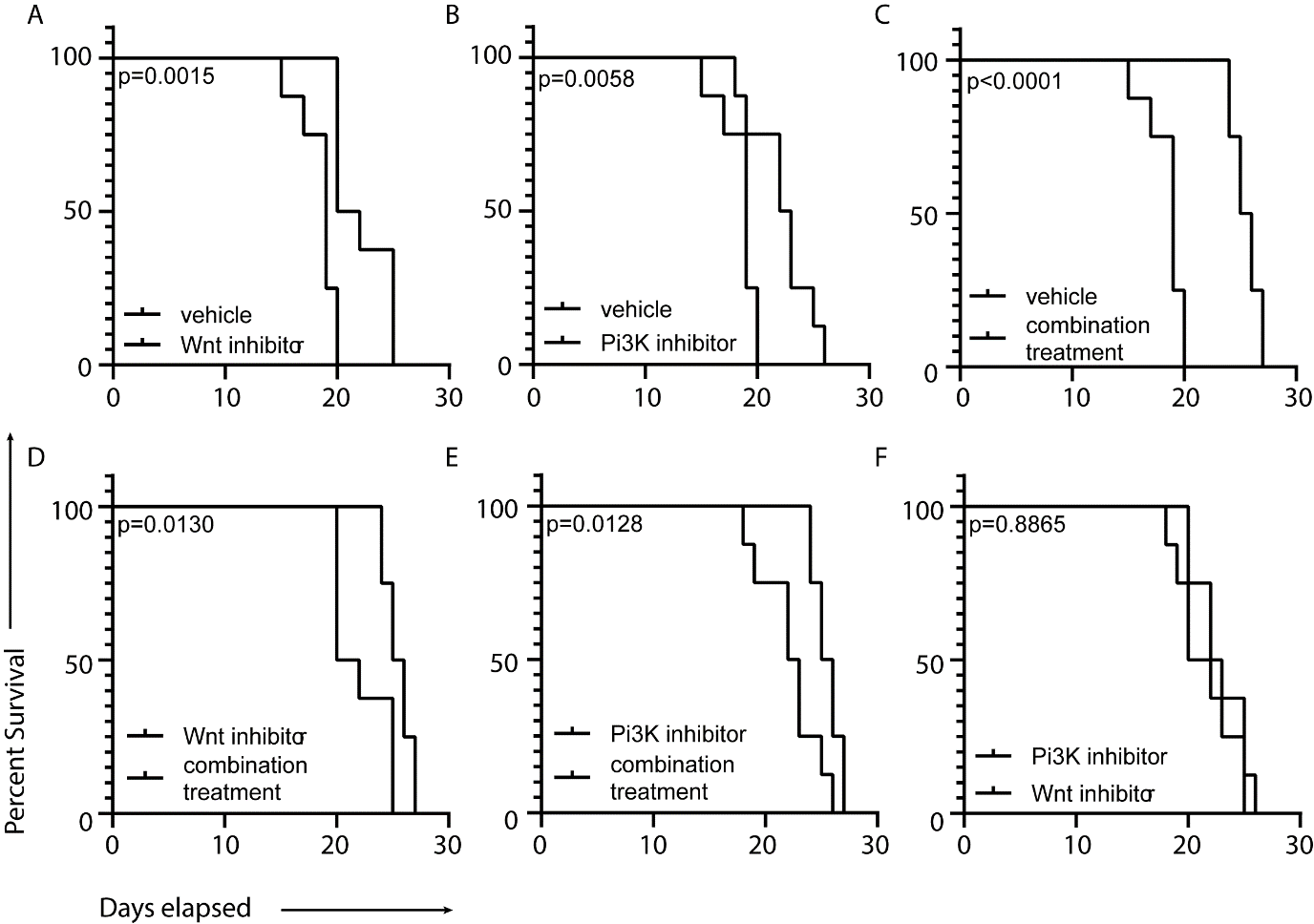

**Supplementary Figure 10 – Segregated survival data for animals treated with Wnt and PI3K inhibitors.** **A-F.** Kaplan-Meier curves of animals baring KRAS^G12D^;*Trp53*^KO^;*Nf2*^KO^ tumours treated with either vehicle, LGK974 (a Wnt inhibitor), Pictilisib (a PI3K inhibitor) or a combination of the two.
